# Metabolic sensor AMPK licenses CD103^+^ dendritic cells to induce Treg responses

**DOI:** 10.1101/2023.02.21.528293

**Authors:** Thiago A Patente, Eline C Brombacher, Graham A Heieis, Leonard R Pelgrom, Anna Zawistowska-Deniziak, Frank Otto, Arifa Ozir-Fazalalikhan, Alwin J van der Ham, Bruno A Guigas, José AM Barbuto, Bart Everts

## Abstract

Dendritic cells (DCs) play a crucial role in promoting tolerance through priming of regulatory T cells (Treg). Several studies indicate DC tolerogenicity is dependent on catabolic metabolism. However, the role of AMP-activated Kinase (AMPK), a key energy and nutrient sensor driving catabolic metabolism, in this process is unclear. We found that human retinoic acid-induced tolerogenic CD103^+^ DCs (RA-DCs) display increased AMPK signaling. Interestingly, RA-DCs, but not vitamin-D3- or dexamethasone-induced tolerogenic DCs, required AMPK for Treg induction. Mechanistically, AMPK underpinned RA-driven tolerogenicity by promoting RALDH activity in a FoxO3-dependent manner. Correspondingly, mice deficient for AMPK in DCs (CD11c^ΔAMPKα1^) harbored reduced frequencies of intestinal CD103^+^CD11b^+^ DCs with impaired RALDH activity. Importantly, upon infection with parasitic worm *Schistosoma mansoni*, that elicits strong Th2 and Treg responses, CD11c^ΔAMPKα1^ mice showed a defect in Treg accumulation and concomitantly, displayed an impaired ability to control Type 2 immunity-driven granulomatous inflammation against the parasite eggs. Together, our findings identify AMPK as a key regulator of tolerance by CD103^+^ DCs.

**Summary:** Dendritic cells (DCs) are critical for inducing tolerance. However, how metabolic cues control their tolerogenicity is still poorly understood. Patente et al demonstrate that AMPK is crucial for Treg induction by retinoic acid-primed tolerogenic CD103^+^ DCs.

## Introduction

Dendritic cells (DCs) are professional antigen-presenting cells that regulate the priming and maintenance of CD4^+^ and CD8^+^ T cell responses and thus play an important role in the development of adaptive immune responses (Steinman and Witmer, 1978). DCs are also key players in maintaining immune tolerance by preventing T cell proliferation and promoting the development of regulatory T cells (Tregs) (Zhao et al., 2018; Steinman et al., 2000). DC populations with tolerogenic properties (tolDCs) commonly express CD103 and can be found in most peripheral tissues, where they play a crucial role in orchestrating peripheral Treg (pTreg) responses and in maintaining tissue-specific tolerance and homeostasis. Most insights in the biology of these tolDCs have been gained from studies on CD103^+^DCs residing in the lamina propria of the gut. Here, acquisition of tolerogenic properties by intestinal DCs is largely driven by dietary Vitamin A, which through its active metabolite RA (Iliev et al., 2009b; a), induces DCs to become CD103^+^ and express retinaldehyde dehydrogenase (RALDH)2 that allows them to produce RA themselves. This RA, in conjunction with TGF-β licenses CD103^+^DCs to prime pTregs (Coombes, 2007; Mucida, 2007). The biology of tolDCs has also been extensively studied in *in vitro* models in which DCs can be rendered tolerogenic following exposure to immunesuppressive or modulatory compounds, including RA, as a model for CD103^+^DCs (Bakdash et al., 2015), dexamethasone (Maggi et al., 2016; Navarro-Barriuso et al., 2018; Jauregui-Amezaga et al., 2015) and the active metabolite of vitamin D3 (Ferreira et al., 2014, 2015; Vanherwegen et al., 2018). Depending on the type of tolerogenic stimulus, different mechanisms have been proposed to control the differentiation of Tregs by these tolDCs. In addition to increased RALDH activity (Kaisar et al., 2017), this includes low costimulatory signal strength (Lan et al., 2006), increased expression of surface bound Immunoglobulin-like transcript members, soluble factors such as IL-10, TGF-β, (Torres-Aguilar et al., 2010), and/or of indoleamine 2,3-dioxygenase (IDO) (Rodrigues et al., 2016; Holtzhausen et al., 2015).

Recently, it has become clear that immune cell activation and function, including that of DCs, are closely linked to changes in cellular metabolism (Buck et al., 2015; Pearce and Everts, 2015; Patente et al., 2019). While the metabolic requirements for the acquisition of an immunogenic phenotype by myeloid DCs are fairly well characterized (Everts et al., 2014, 2012; Guak et al., 2018), the metabolic pathways involved in the development and function of tolDCs, are less well defined (Malinarich et al., 2015; Zhao et al., 2018). There is some evidence that tolDCs rely on oxidative phosphoryalation (OXPHOS) for their tolerogenic functions (Malinarich et al., 2015; Zhao et al., 2018) and it has been shown that inhibiting anabolic metabolism in DCs promotes the development of a tolerogenic phenotype (Fischer et al., 2009), indicating a role for catabolic/oxidative metabolism in Treg induction by DCs. However, currently, how exogenous tolerogenic signals are integrated to promote this catabolic profile in tolDCs are still largely unknown.

AMP-activated protein kinase (AMPK) is a key sensor of cellular energy status, restoring energy homeostasis upon nutrient deprivation by inhibiting anabolic pathways and promoting catabolic pathways (González et al., 2020; Steinberg and Hardie, 2022). There is some data to suggest a role for AMPK in shaping the immunogenic properties of DCs. For instance, human DCs treated with VitD3 displayed increased AMPK activation, which was associated with increased mitochondrial activity (Ferreira et al., 2015) and elevated mitochondrial glucose oxidation (Vanherwegen et al., 2018), suggesting a possible role for AMPK signaling in tolDCs differentiation via metabolic alterations. In addition, AMPK deficiency in CD11c-expressing cells was shown to hamper host defense against hookworm infection in mice, a feature associated with increased IL-12/23p40 secretion and poor induction of a type 2 immune response (Nieves et al., 2016). Concomitantly, murine bone marrow-derived DCs (BMDCs) from AMPK knockout mice displayed an increased inflammatory phenotype characterized by an enhanced ability to induce IFN-γ- and IL-17-secreting T cells (Carroll et al., 2013). While these studies suggest that AMPK signaling in DCs is involved in dampening immune responses, the exact role played by this kinase in tolDC induction, metabolism and function remains an open question.

Here, we set out to characterize the metabolic properties of differently generated tolDCs and to define the role of AMPK signaling in their differentiation and ability to induce Tregs. We show that specifically human retinoic acid-induced tolerogenic CD103^+^ DCs (RA-DCs) are characterized by increased AMPK signaling and that this AMPK signaling is required for the induction of functional Tregs. We found AMPK to mediate this via increasing RALDH activity, through FoxO3-signaling, independently from metabolic rewiring. Similarly, murine intestinal tolerogenic RALDH^+^CD103^+^ DCs displayed high AMPK activation and their RALDH activity was abrogated upon loss of AMPK signaling. Furthermore, we link the loss of AMPK in DCs to impaired Treg induction and exacerbated pathology during infection with the parasitic helminth *Schistosoma mansoni*. Taken together, these data identify a key role for AMPK signaling in RA-driven development of tolDCs, that mitigate tissue pathology during inflammation.

## Results

### Differently generated human tolDCs display distinct metabolic profiles

To characterize and compare the metabolic properties of different tolDCs, we generated moDCs from peripheral blood CD14^+^ monocytes with GM-CSF and IL-4, followed by LPS stimulation in the absence (mDC) or presence of three well-known tolerogenic compounds: retinoic acid (RA-DCs) (Bakdash et al., 2015), dexamethasone (Dex-DCs) (Maggi et al., 2016; Navarro-Barriuso et al., 2018; Jauregui-Amezaga et al., 2015) and the active metabolite of vitamin D3 (VitD3-DCs) (Ferreira et al., 2014, 2015; Vanherwegen et al., 2018). In accordance with previous studies (Ferreira et al., 2013; Does et al., 2014), all tolerogenic compounds reduced moDC differentiation, as indicated by reduced CD1a expression and increased CD14 expression in the case of VitD3 **(Figure S1A).** Furthermore, in line with literature (Unger et al., 2009; Penna et al., 2005; Manavalan et al., 2003) VitD3, but not Dex, significantly increased the expression of PD-L1 and ILT3, two co-inhibitory molecules involved in controlling the inflammatory response (Chang et al., 2009), while RA treatment increased the expression of the gut homing integrin, CD103 **(Figure 1A and Figure S1B)** as well as RALDH activity **(Figure 1B).** The LPS-induced costimulatory molecule expression by DCs was overall inhibited when cells were treated with VitD3, RA and Dex **(Figure 1A),** but only VitD3 and Dex reduced the secretion of IL-12p70 and increased the secretion of IL-10 by DCs **(Figure 1C).** Importantly, irrespective of these phenotypic differences, all tolDCs induced stable and functional Tregs capable of suppressing bystander T cell proliferation **(Figure 1D)** and that are characterized either by increased CTLA-4 (by all tolDCs; **Figure S1C)** Foxp3 (by RA-DCs; **Figure 1E)** or IL-10 expression (by Dex- and RA-DCs; **Figure 1F).**

**Figure 1:**
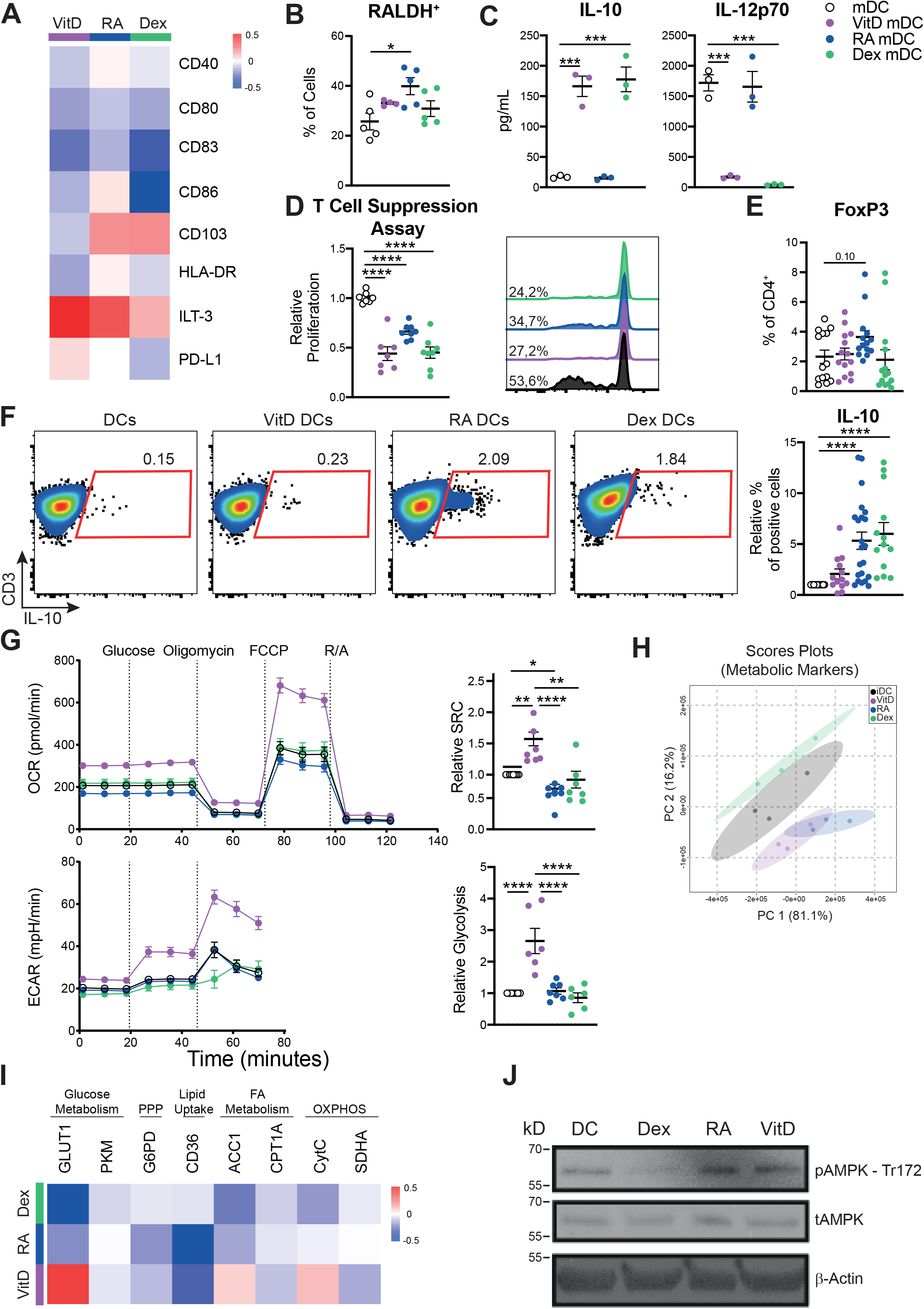
Differently generated human tolerogenic DCs display distinct metabolic profiles. Monocytes were isolated from PBMCs with CD14^+^ magnetic beads and differentiated in moDCs in the presence of GM-CSF + IL-4. On day 5 cells were treated with either VitD3, RA or Dex. On the 6^th^ day, 100 ng/mL of LPS were added and 24 hours later cells were harvested for functional assays. **(A)** Heatmap of the log2 fold change in geometric mean fluorescence intensity (GeoMFI) of surface markers of tolDCs measured by flow cytometry relative to control-stimulated DCs (mDC) (set to 0). **(B)** frequency of RALDH^+^ cells in tolDCs measured by flow cytometry. **(C)** tolDCs were co-cultured with CD40L-expressing cell line (J558) for 24 hours, supernatants were collected, and cytokines were measured by ELISA. **(D)** A T cell suppression assay was performed by co-culturing irradiated TolDC primed naïve Th cells with CFSE-labelled memory Th cells from the same donor. On the 6^th^ day CFSE dilution was measured by flow cytometry. (E-F) tolDCs were co-cultured with allogeneic naïve T cells and intracellular **(E)** FoxP3 staining and **(F)** levels of IL-10 were measured by flow cytometry. **(G)** Seahorse extracellular flux analysis. Real-time oxygen consumption rate (OCR) (upper left) and extracellular acidification rate (ECAR) (bottom left) in tolDCs. Spare respiratory capacity (SRC) (upper right) and glycolysis (bottom right) were quantified and are displayed as fold change compared to control-stimulated DCs (mDC). (H) Principal component analysis (PCA) of tolDCs determined by the expression pattern of metabolic proteins. (I) Heatmap of the normalized geoMFI values of intracellular staining for metabolic proteins as determined by flow cytometry relative to mDCs. (J) Representative immunoblot of the expression of phosphorylated AMPK, total AMPK and β-actin in DC either left alone or treated with Dex, RA or VitD3 for 48 hours. Data are pooled from 15-22 donors **(A, E, F),** 5-7 donors **(B, D, G)** or 3 donors **(H,I),** or are representative of 1 out of 3 independent experiments (C), or 2 independent experiments (J). Mean ± SEM are indicated in the graphs. One-way Anova with Tukey HSD post-test was used to assess statistically significant differences; *p < 0.05, **p < 0.01, ***p < 0.001, ****p<0.0001.

Following validation of their tolerogenic phenotype and function we next investigated the metabolic profile of the differently generated tolDCs. VitD3 induced the most pronounced core metabolic alterations, as evidenced by increased rates in glycolysis and OXPHOS **(Figure 1G).** On the other hand, Dex did not induce any core metabolic changes, while RA only reduced the spare respiratory capacity (SRC) of DCs **(Figure 1G).** In contrast to control DCs, all tolDCs showed increased levels of OXPHOS upon LPS stimulation **(Figure S2),** which is indicative of increased mitochondrial glucose oxidation and catabolic metabolism. Flow cytometry-based analysis of expression of rate-limiting metabolic enzymes in key metabolic pathways (Heieis et al., 2022; Pelgrom et al., 2022), corroborated these findings, by revealing that amongst the different tolDCs, VitD3-DCs displayed the most pronounced changes in metabolic enzyme expression **(Figure 1I, S1D).** Although principal component analysis of the metabolic enzyme expression data showed co-clustering of VitD3-DCs with RA-DCs, but not Dex-DCs **(Figure 1H),** modulation of several metabolic enzymes by RA was distinct from VitD3, including reduction in Acetyl-CoA Carboxylase (ACC)1 and glucose transporter (Glut)1 expression, potentially indicating a more catabolic metabolic profile **(Figure 1I).**

Finally, we assessed activation status of AMP-activated protein kinase (AMPK) as one of the central energy sensors controlling catabolic metabolism. Interestingly increased phosphorylation of AMPK at Thr172 **(Figure 1J),** as well as of its direct substrate, ACC at Ser79 **(Figure S1E),** was observed in both VitD3- and RA-DCs, but not Dex-DCs. Together these data reveal that different tolDCs with distinct tolerogenic immune features also display divergent metabolic programs as well as levels in AMPK signaling.

### RA-DCs depend on AMPK signaling for induction of regulatory T cells

In light of the known link between tolDC function and catabolic metabolism, we next aimed to explore the functional relevance of high AMPK signaling in the differently generated tolDCs. To this end, we silenced AMPKα1, the primary isoform expressed by DCs (Nurbaeva et al., 2012), prior to treatment with the tolerogenic compounds **(Figure S3A)** and evaluated whether the lack of AMPKα1 signaling would impair phenotype and function of the differently stimulated tolDCs. We found that the metabolic alterations induced by in VitD3 and RA compounds were reversed in the absence of AMPKα1 but not in Dex-DCs **(Figure 2A, 2B and S3B).** Next, we investigated the effects of AMPKα1 signaling on the ability of tolDCs to induce the differentiation of *bona-fide* Tregs and found that AMPKα1 silencing did not impair Dex-induced tolerogenicity **(Figure 2C).** Surprisingly, despite the reversal of metabolic alterations induced by VitD3 following AMPKα1 silencing, AMPKα1 signaling was dispensable for the induction of functional Tregs by VitD-DCs **(Figure 2C).** Importantly, on the other hand, lack of AMPKα1 significantly compromised the ability of RA-DCs to drive IL-10 production in T cells and to promote their differentiation into functional Tregs **(Figure 2C, 2D).** To further substantiate these findings, we generated murine RA-BMDCs from mice with a specific deletion of AMPKα1 in CD11c^+^ positive cells (CD11c^ΔAMPKa1^) **(Figure S3C)** and assessed their ability to promote differentiation of FoxP3^+^ Tregs and to suppress OT-II T cell proliferation. In line with the human DC data, AMPKα1-deficient RA-BMDCs failed to upregulate Foxp3 in cocultured CD4^+^ T cells and to suppress T cell proliferation, in contrast to their WT counterparts **(Figure 2E, 2F).** Together, the data show that VitD3 and RA increase AMPKα1 signaling in DCs and that, while AMPKα1 is important for the metabolic alterations observed by both tolerogenic compounds, only RA-DCs rely on AMPKα1 for their tolerogenic functions.

**Figure 2:**
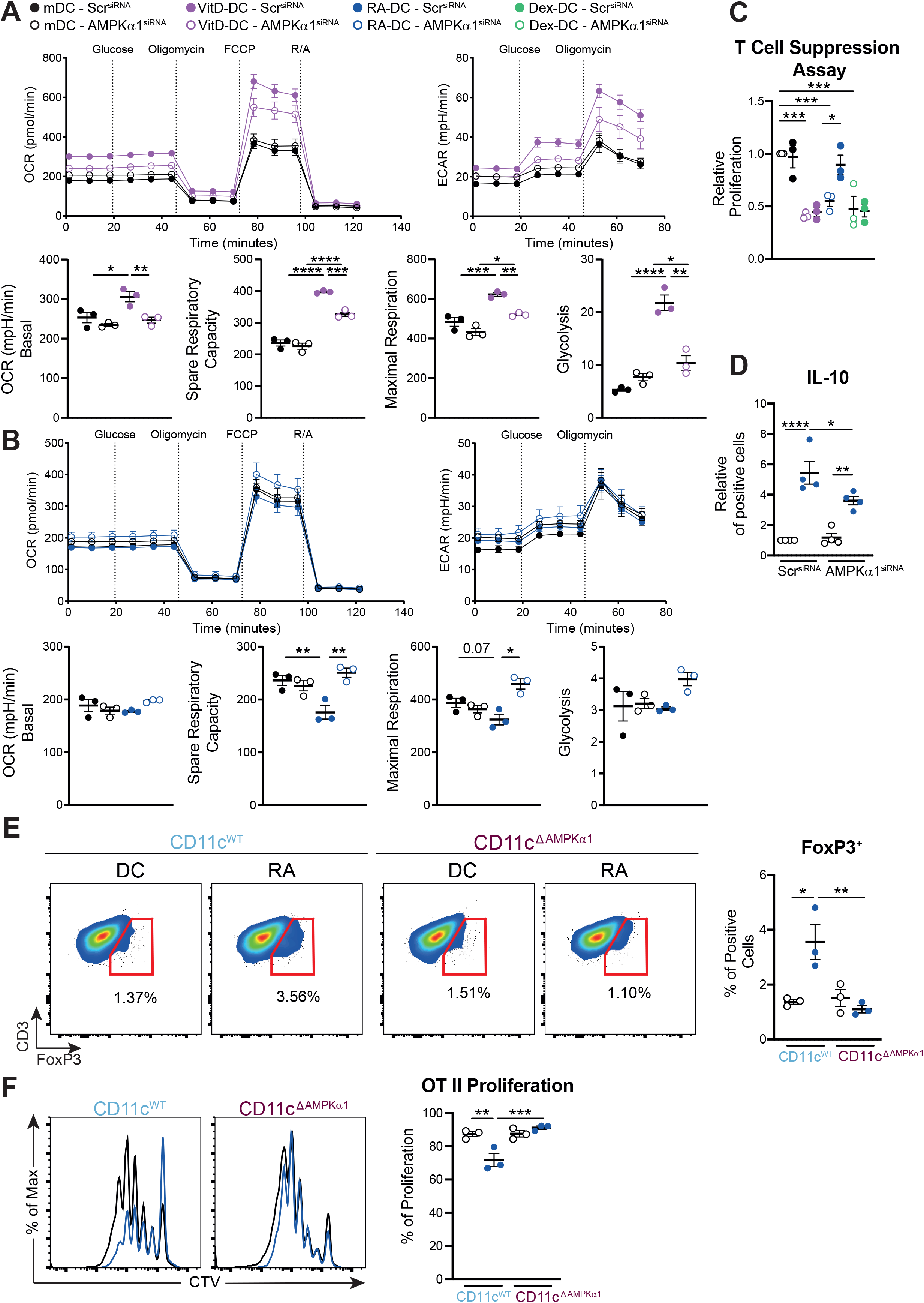
RA-DCs depend on AMPK signaling for induction of regulatory T cells. moDCs were silenced for AMPKα1 or non-targeting scramble RNA before treatment with VitD3 or RA. Subsequently, metabolic profile for VitD- **(A)** and RA-treated DCs **(B)** was analyzed using Seahorse extracellular flux analysis and suppressive capacity **(C)** and IL-10 secretion **(D)** of T cells primed by tolDCs were assessed. **(E-F)** GM-CSF-derived bone marrow dendritic cells (GMDCs) were differentiated from CD11c^WT^ orCD11c^ΔAMPKα1^ mice and co-cultured with OT II naïve T cells to evaluate their ability to induce **(E)** FoxP3^+^ Tregs or **(F)** their ability to induce OT II T cell proliferation. Data are representative of 1 out of 4 donors (A-B), represent 3-4 donors (C-D), or 3 independent experiments **(E-F).** Mean ± SEM are indicated in the graphs. Two-Way Anova with Sidák’s post-test was used to assess statistically significant differences.; *p < 0.05, **p < 0.01, ***p < 0.001, ****p<0.0001.

### AMPK licenses RA-DCs to promote Tregs through induction of RALDH activity

In the light of these results, we next aimed to address through which mechanisms AMPKα1 signaling renders RA-DCs tolerogenic. To this end, we assessed the consequences of AMPKα1 silencing on the immunological phenotype of RA-DCs. We found AMPKα1 signaling to be important for the reduction of CD80 and for the increase in both CD103 expression and RALDH activity induced by RA **(Figure 3A, 3B and Figure S4A).** RA-induced RALDH activity in DCs has been shown to be important for the induction of IL-10-secreting T cells and promotion of functional Tregs (Bakdash et al., 2015; Feng et al., 2010; Kaisar et al., 2017). Indeed, following treatment with RALDH inhibitor diethylaminobenzaldehyde (DEAB), prior to RA treatment, the ability of human RA-DCs to promote functional Tregs, **(Figure 3C)** or of murine RA-DCs to suppress proliferation of OT II T cells was lost **(Figure S4B).** This suggests that AMPKα1 signaling underpins the tolerogenic function of RA-DCs through induction of RALDH activity.

**Figure 3:**
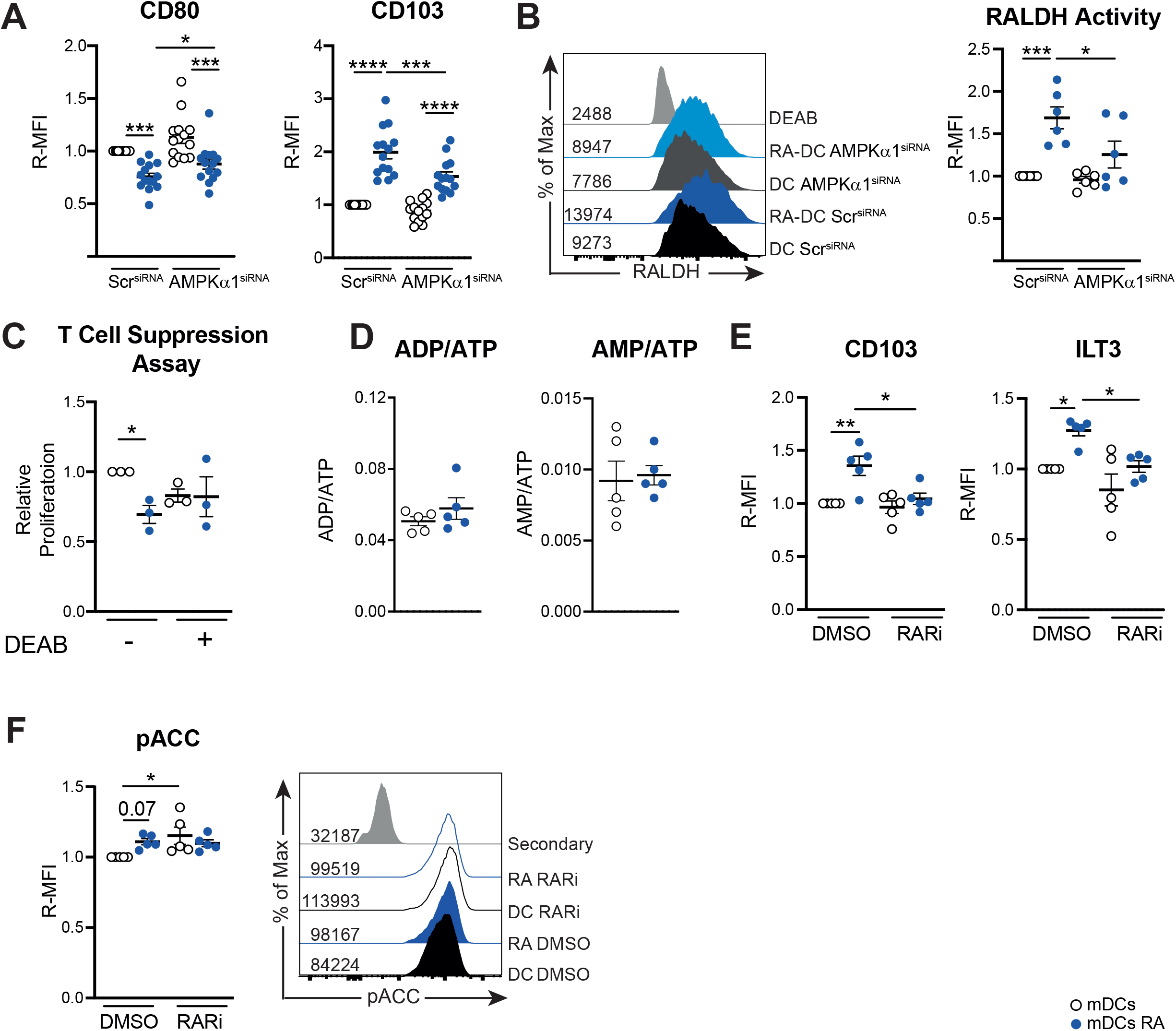
AMPK licenses RA-DCs to promote Tregs through induction of RALDH activity. **(A)** Relative geoMFI (R-MFI) for the expression of CD80 and of CD103 and from AMPKα1-silenced or scramble-silenced RA-DCs measured by flow cytometry. **(B)** Representative histogram staining for RALDH activity (left) and quantification of R-MFI in RA-DCs silenced for AMPKα1. **(C)** moDCs were pre-treated with either vehicle or DEAB for 30 min before being treated with RA followed by LPS stimulation and T cell suppression assay was performed as described in Figure 1D. **(D)** Intracellular levels of AMP, ADP and ATP in moDCs were measured by HPLC and the ratio of ADP/ATP and AMP/ATP were quantified. (E-F) moDCs were pre-treated with either vehicle or retinoic acid receptor inhibitor (RARi) for 1 hour before being treated with RA and **(E)** relative geoMFI (R-MFI) for the expression of CD103 and ILT-3 were evaluated. **(F)** Intracellular levels of phospho acetyl carboxylase (serine 79 - pACC) levels were evaluated in moDCs pre-treated with RARi followed by 4 h of RA treatment (left); representative histogram of the pACC staining on moDCs (right). Data are based on a pool of 14 **(A)**; 6 **(B)**, or 5 donors **(D-F)**. Mean ± SEM are indicated in the graphs. Two-Way Anova with Sidák’s post-test **(B-C;E-F)** or student’s T test **(D)** were used to assess statistically significant differences.; *p < 0.05, **p < 0.01, ***p < 0.001.

To understand how RA stimulates AMPKα1 signaling in moDCs, we first evaluated whether RA treatment could affect cellular energy status by measuring the intracellular levels of AMP, ADP and ATP, as increased ratios in AMP/ATP and ADP/ATP promote AMPKα1 activation (Steinberg and Hardie, 2022; González et al., 2020). However, RA neither alter ATP/ADP nor AMP/ATP ratios, suggesting RA does not increase AMPKα1 activity through changes in bioenergetic status **(Figure 3D).** We next explored whether RA may affect AMPKα1 signaling through binding with its nuclear receptor, retinoic acid receptor (RAR). While preincubation with pan RAR inhibitor AGN 194310 prevented RA-induced changes in expression of CD103 and ILT3 **(Figure 3E),** it did not affect the increase in AMPKα1 signaling induced by RA treatment **(Figure 3F).** Together, these data suggest that RA in DCs, independently from changes in bioenergetics or RAR signaling, promotes AMPKα1 activation which is required for increased RALDH activity and thereby induction of Tregs by RA-DCs.

### AMPKα1 signals via PGC1α to control CD103 expression and via FoxO3 to control RALDH activity

We next set out to investigate the mechanism through which AMPKα1 drives RALDH activity in DCs. PGC1α is a substrate of AMPK (Jäger et al., 2007) that can act as a co-activator for the nuclear receptor PPARγ (Liang and Ward, 2006). As it has been proposed that PGC1α controls RALDH activity in murine CD103^+^CD11b^-^ DCs (Jin et al., 2020), we wondered whether AMPKα1-driven RALDH activity in RA-DCs is dependent on PPARγ-PGC1α signaling. We found mRNA expression of both PPARγ and PGC1α to be significantly upregulated in response to RA treatment in DCs **(Figure 4A).** However, although pharmacological inhibition of PGC1α did prevent RA-induced expression of CD103 **(Figure 4B),** neither inhibition of PGC1α nor PPARγ did affect the ability of RA to increase RALDH activity **(Figure 4C)** nor of RA-DCs to prime Tregs **(Figure S4C),** suggesting that AMPKα1 does not signal through the PPARγ-PGC1α to induce RALDH activity and tolerogenicity in RA-DCs. Another transcription factor downstream of AMPK is FoxO3, which belongs to the forkhead-box family of transcription factors and plays a role in a variety of processes including longevity, cell differentiation and energy homeostasis (Fasano et al., 2019). *in silico* analysis revealed that the top phosphorylated site by AMPK in FoxO3 is S413, which has been shown to promote its transcriptional activity (Greer et al., 2007) **(Figure 4D).** Congruently, we found that upon RA treatment, DCs displayed increased phosphorylation at S413, **(Figure 4E).** In addition, phosphorylation of FoxO3 at S253 which is known to prevent its translocation to the nucleus (Brunet et al., 1999), was inhibited **(Figure 4E).** Importantly, silencing AMPKα1 in moDCs abolished the RA-induced phosphorylation of FoxO3 at S413 **(Figure 4F).** In addition, while silencing of FoxO3 in moDCs did not affect expression of surface markers modulated by RA, including CD103 **(Figure 4G, S4D),** the ability of RA to induce RALDH activity and promote tolerogenic DC function was lost **(Figure 4H-I).** Taken together, these data show that RA triggers AMPKα1 signaling to promote RALDH activity in a FoxO3-dependent manner that is required for subsequent Treg induction.

**Figure 4:**
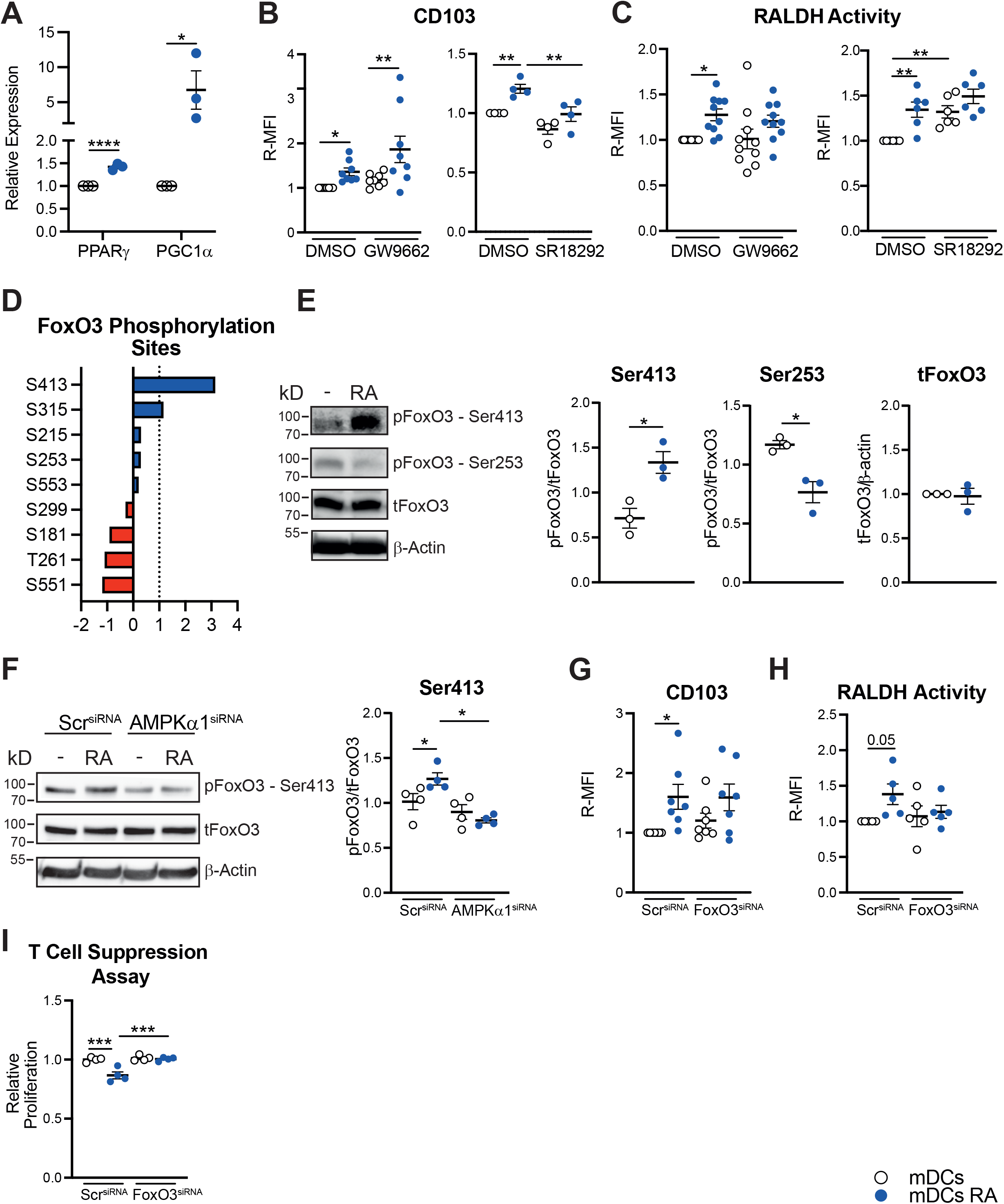
AMPK signals via PGC1α to control CD103 expression and via FoxO3 to control RALDH activity. **(A)** Gene expression of PPARγ and PGC1α in RA-treated and control moDCs. On day 5 of culture moDCs were treated either with RA or left untreated for 48 hours and mRNA expression levels were determined by RT-qPCR. GAPDH was used as housekeeping gene **(B-C)**. On day 5, moDCs were pre-treated with either DMSO, PPARγ (GW9662) or PGC1α (SR18292) inhibitors for 30 minutes prior to RA treatment, followed by LPS stimulation and **(B)** CD103 expression and **(C)** RALDH activity were quantified by flow cytometry. **(D)** *In silico* analysis displaying the top predicted AMPK-phosphorylation sites in FoxO3. Dashed line represents the threshold level of AMPK-FoxO3 interaction. **(E)** Representative immunoblot (left) and quantification (right) of the expression levels of FoxO3 Ser413, FoxO3 Ser253, total FoxO3 and β-actin in RA-DCs. **(F)** Representative immunoblot (left) and quantification (right) of the expression levels of FoxO3 Ser413, total FoxO3 and β -actin in RA-DCs silenced for AMPKa1 as described in Figure 2. **(G)** Expression levels of CD103 and **(H)** RALDH activity in RA-DCs silenced for FoxO3 as described in Figure 2. **(I)** moDCs were silenced for FoxO3 as described in Figure 2 and T cell suppression assay was performed as described in Figure 1D. Data are a pool of 3 **(A)**; 4-8 **(B-C)**, 3-4 **(E-F)**, or 4-7 donors (G-H) or are representative for 1 out of 3 independent experiments **(I)**. Statistics are student’s T-test **(A;E)** and Two-Way Anova with Sidák’s posttest **(B-C;F-I)**, were used to assess statistically significant differences. Mean ± SEM are indicated in the graphs; *p < 0.05, **p < 0.01, ***p < 0.001, ****p<0.0001.

### AMPKα1 signaling governs maintenance and RALDH activity of intestinal CD103^+^ CD11b^+^ DCs

Next, we explored the functional relevance of these findings in an *in vivo* setting. Naturally occurring tolerogenic RALDH^+^CD103^+^ DCs have been identified in various organs, including lung and intestine (Guilliams et al., 2010; Ruane et al., 2013). We initially focused on the role of AMPK signalling in the biology of RALDH^+^CD103^+^ DCs in the latter organ, since here DCs are thought to be exposed to particularly high levels of RA readily generated by epithelial cells from dietary vitamin A (Iliev et al., 2009b; a; Mucida et al., 2007; Coombes et al., 2007). In the intestine, three main conventional DC subsets can be identified, based on differential expression of CD103 and CD11b **(Figure S5A)** of which particularly CD103^+^ DCs populations are associated with the induction of Tregs (Esterházy et al., 2019). We observed that in naïve WT mice small intestinal lamina propria (siLP) CD103^+^ DCs (especially CD103^+^CD11b^+^) displayed higher levels of pACC compared to CD11b^+^ DCs, CD64^+^ macrophages (Mph), T cells or B cells from the same tissues **(Figure 5A)** as well as compared to DCs from other tissues including lung, liver, spleen and msLN **(Figure S6A, S6B).** Interestingly, AMPK signaling levels were positively correlated with RALDH activity in these DCs **(Figure 5A).** Largely in line with our *in vitro* data, we found that mice with a specific deletion of AMPKα1 in CD11c^+^ cells (CD11c^ΔAMPKα1^) **(Figure S6C),** showed a selective reduction in cell frequency of CD103^+^CD11b^+^ DCs from siLP **(Figure 5B)** as well as a lower RALDH activity in this DC subset **(Figure 5C).** Reduced frequencies of CD103^+^CD11b^+^ DCs in CD11c^ΔAMPKα1^ mice were mirrored in the draining mesenteric (mes)LNs **(Figure S6F),** although RALDH activity was not affected in these DCs by loss of AMPKα1 **(Figure S6G).** To further characterize the consequences of loss of AMPKα1 on siLP DC populations, we performed an unbiased analysis of the intestinal DC compartment, through dimensional reduction and unsupervised analysis based on a flow cytometry panel encompassing activation and lineage markers. This resulted in the identification of several different phenotypic clusters within the three subsets of intestinal DCs **(Figure 5D, 5E and S6D).** Amongst those, cluster PG_04, a CD103^+^CD11b^+^ population characterized by high RALDH activity and low CD86 and CCR7 expression, was significantly reduced in frequency in CD11c^ΔAMPKα1^ mice, while the opposite was true for a RALDH^lo^CD86^hi^CCR7^hi^ CD103^+^ DC population (PG_09) **(Figure 5F, 5H and S6E),** indicative of a switch from a tolerogenic to a more immunogenic intestinal DCs phenotype in CD11c^ΔAMPKα1^ mice.

**Figure 5:**
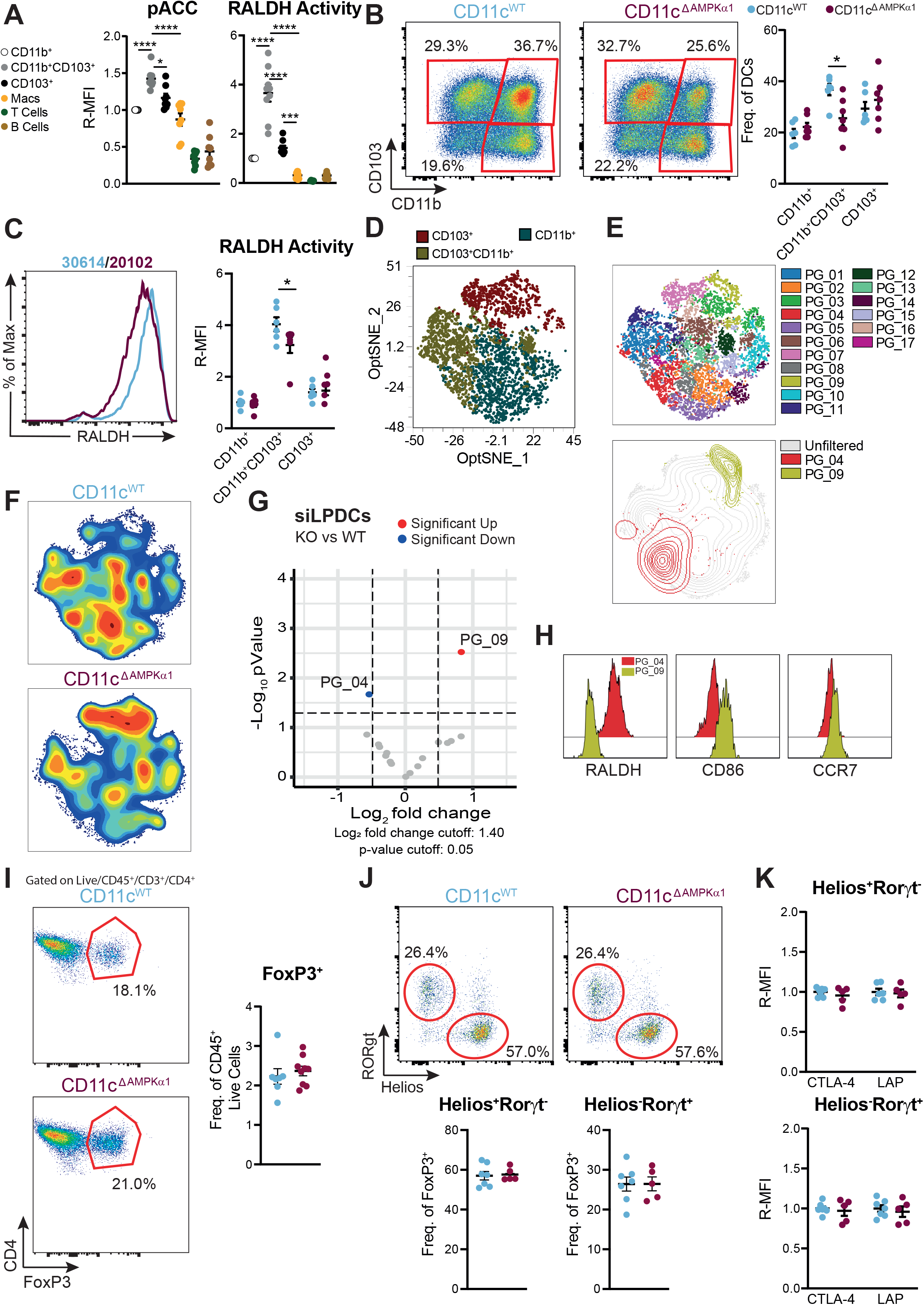
AMPK signaling governs maintenance of, and RALDH activity in intestinal CD103^+^CD11b^+^ DCs. **(A)** Relative geometric MFI (R-MFI) of phosphorylation level of acetyl-coA carboxylase -Ser79 (pACC - left) and retinaldehyde dehydrogenase (RALDH) activity in different immune cells in the small intestine lamina propria (siLP) of naïve WT mice. Data were normalized by the expression levels of CD11b^+^ DCs. **(B)** Representative dot plot (left) and frequency of siLP DC subsets within CD45^+^ live cells (right) in CD11c^WT^ (salem blue) and CD11c^ΔAMPKα1^ (purple-red). **(C)** Representative histogram (left) and relative geometric MFI (R-MFI) of RALDH activity in siLP DC subsets. **(D)** Unbiased opt-SNE analysis of siLP DC populations in which CD103^+^, CD103^+^CD11b^+^ and CD11b^+^ are indicated. **(E)** Phenograph clustering performed on siLP DCs using RALDH, activation and lineage markers. **(F)** Contour plots overlaid on opt-SNE analysis as shown in **(D)** displaying distribution of cells for CD11c^WT^ and CD11c^ΔAMPKα1^ mice. **(G)** Volcano plot displaying the clusters as displayed in **(E)** with significant differences in frequency between CD11c^WT^ and CDllc^ΔAMPKα1^ mice. **(H)** Histograms displaying the expression levels of RALDH, CD86 and CCR7 in PG_04 (red) and PG_09 (ochre). **(I)** Representative dot plot for the FoxP3^+^ regulatory T cells (Tregs) in the siLP of CD11c^WT^ and CD11c^ΔAMPKα1^ mice (left) and frequency of Tregs within CD45^+^ live immune cells (right). **(J)** Representative dot plot for the thymic (Helios^+^RORgt^-^ - tTregs) and peripheral (Helios^-^RORgt^+^ - pTregs)) Tregs in the siLP of CD11c^WT^ and CD11c^ΔAMPKα1^ mice (top) and frequency of tTregs and pTregs within total Treg population (bottom). **(K)** Quantification of CTLA-4 and latency-associated peptide (LAP) as proxy of TGF-b secretion by tTregs and pTregs in CD11c^WT^ and CD11c^ΔAMPKα1^ mice. Data are representative of 1 out of 3 (left) or a pool of 3 independent experiments **(A)**; represent a pool of 2 to 3 independent experiments with 3-4 mice per experiment **(A-C;I-K)**; representative of 1 out of 2 independent experiments **(D-H).** One-Way Anova with Tukey HSD post-test **(A)** or student’s T-test **(B-C-K)** were used to assess statistically significant differences. Mean ± SEM are indicated in the graphs; *p < 0.05, ***p < 0.001, ****p<0.0001.

We also analyzed the lung as another mucosal site with regulatory characteristics where RALDH^+^CD103^+^ DCs have been identified (Guilliams et al., 2010). We did not observe any significant changes in frequencies and RALDH activity at the population level in CD11b^+^ or CD103^+^ DC subsets in the lungs of CD11c^ΔAMPKα1^ compared to CD11c^WT^ mice **(Figure S7A-C).** However, when analyzing the lung DC compartment in an unbiased manner, we identified several phenotypic clusters within the CD103^+^ and CD11b^+^ DC subsets with different abundance between CD11c^WT^ and CD11c^ΔAMPKα1^ mice **(Figure S7D, S7E, and S7F).** Similar to our findings in the gut, we observed a shift from a tolerogenic to an immunogenic DC phenotype in CD11c^ΔAMPKα1^ mice, as illustrated by, on the one hand, a reduced frequency of a CD103^+^ DC cluster, with high RALDH activity and low expression of CD80 and CD86 (PG_04), and, on the other hand, an accumulation of a RALDH^lo^CD86^hi^CD80^hi^ CD103^+^ DC population (PG_07) **(Figure S7G-S7J).**

Despite the differences observed in the CD103^+^ DC compartment in the absence of DC-intrinsic AMPK, neither the frequency nor the activation state of Tregs was significantly affected in lung or intestine of naïve CD11c^ΔAMPKα1^ mice **(Figure 5I-K, S6H, I, S7K).** Analysis of the Treg populations further revealed no differences in the frequency of Tregs from either thymic (tTreg; Helios^+^RORγt^-^) or peripheral origin (pTreg – Helios^-^RORγt^+^) **(Figure 5J, S6H).** Taken together these data suggest that AMPKα1 signaling controls CD103^+^ DCs tolerogenic phenotype by governing RALDH activity, but that this does not grossly affect mucosal Treg populations at steady state.

### AMPK signaling in CD11c-expressing cells is important for Treg induction and prevention of immunopathology during *S. mansoni* infection

Although the Treg compartment of CD11c^ΔAMPKa1^ mice appeared unaffected during steady state, we next wondered whether the altered phenotype of CD103^+^ DCs may affect Treg responses in the context of inflammation. Exposure to house dust mite (HDM) to drive allergic asthma, promoted accumulation of Tregs in the medLNs in WT mice. Interestingly, this was significantly compromised in CD11c^ΔAMPKα1^ mice **(Figure S7L).** However, no differences in cellular infiltrate or eosinophilia in bronchioalveolar lavage **(Figure S7M)** or lungs **(Figure S7N)** were observed, suggesting that the defect in Treg accumulation in the lung, as a consequence of allergen challenge, did not significantly impact the inflammatory response and outcome of disease in CD11c^ΔAMPKα1^ mice.

We next aimed to investigate whether CD11c-specific AMPKα1 loss affected Treg responses and pathology during infection with parasitic helminth *Schistosoma mansoni*. During schistosomiasis major drivers of pathology are the parasitic eggs released by mature worms residing in the mesenteric vasculature that, as a consequence of egg-induced Type 2 inflammation, lead to granulomatous tissue lesions in the intestine but particularly the liver. During this infection induction of pTregs are crucial for preventing overzealous Th2 responses and uncontrolled granuloma formation especially during the chronic stages of infection. We observed the appearance of a CD103^+^CD11b^+^ DC subset in the liver of both acutely (8 week) and chronically (16 week) infected mice **(Figure 6A and S7O)** displaying increased AMPK signaling, as determined by ACC phosphorylation levels **(Figure 6B and S7P),** as well as increased RALDH activity **(Figure 6C and S7Q).** Importantly, in contrast to the WT littermates, infected CD11c^ΔAMPKα1^ mice failed to increase Treg frequencies in their livers at both 8 **(Figure 6D, E)** and 16 weeks post infection **(Figure 6F).** At both timepoints, this was associated with an increase in both Th1 and Th2 cytokine secretion in the hepatic lymph nodes (hpLN) **(Figure 6G, H)** and msLN **(Figure 6I)** of CD11c^ΔAMPKα1^ mice. Consistent with an uncontrolled type 2 immune response, we found a higher frequency of alternatively activated CD11c^-^RELMα^+^ macrophages **(Figure 6J)** as well as an increased liver egg granuloma size in chronically, infected CD11c^ΔAMPKα1^ mice compared to infected WT littermates **(Figure 6K),** indicative of enhanced immunopathology, which was not yet evident during acute stages of infection **(Figure S7R).** Taken together, the data suggest that AMPKα1 signaling in DCs is important to induce the differentiation and or expansion of FoxP3^+^ Tregs, to keep effector Th cell responses overzealous egg-induced granulomatous responses in check during *S. mansoni* infection.

**Figure 6:**
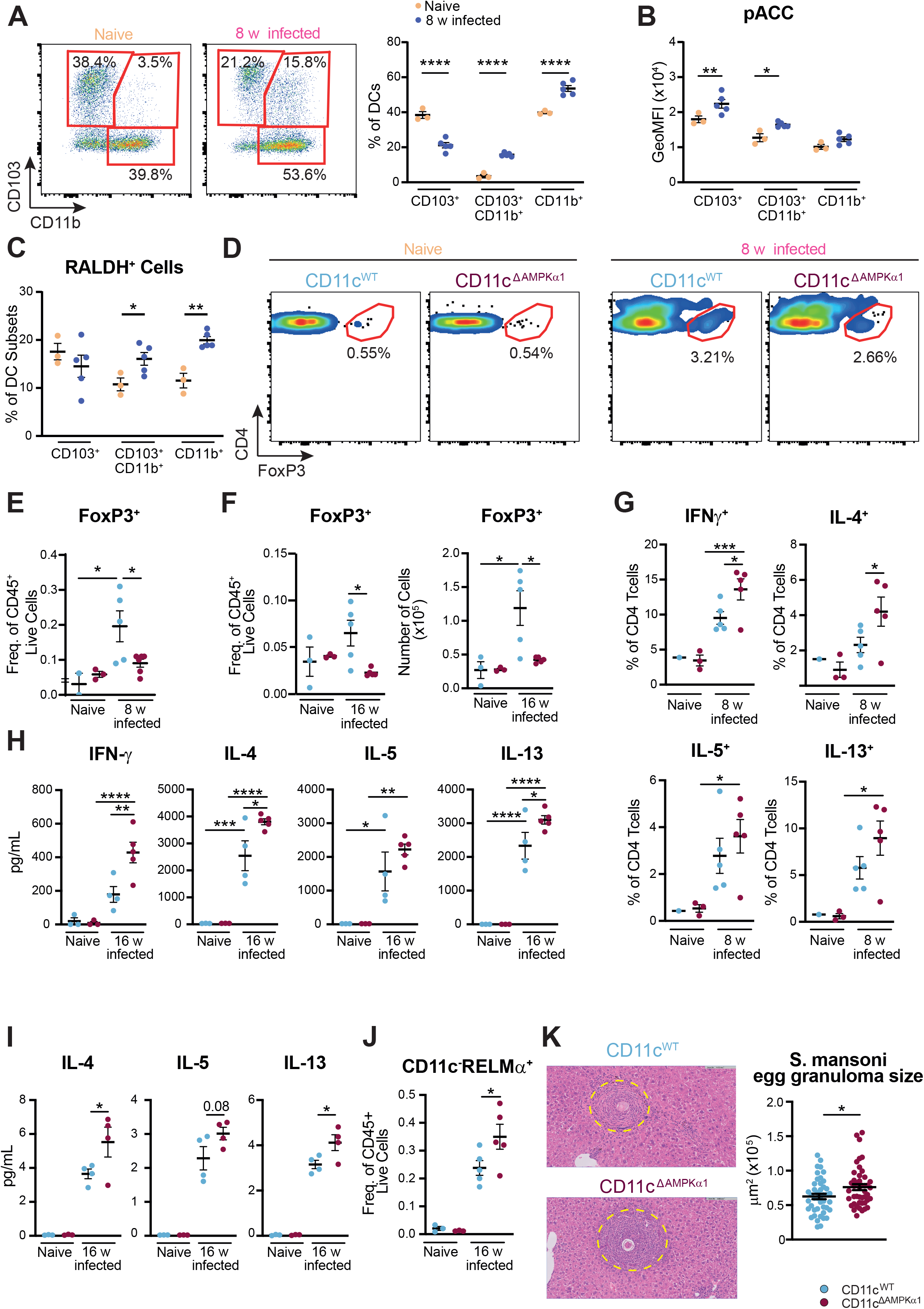
AMPK signaling in CD11c-expressing cells is important for Treg induction and to keep the effector T cell and the granulomatous response in check during *S. mansoni* infection. Mice were infected with *S. mansoni* by percutaneous exposure to cercariae and immune cell responses were evaluated in livers and draining hepatic and mesenteric lymph nodes 8 or 16 weeks later. **(A)** Representative dot plot (left) and frequency of DCs (right) in the liver of naive (yellow) and 8 week-infected WT mice (blue). **(B)** GeoMFI of phosphorylation level of acetyl-coA carboxylase-Ser79 (pACC) and **(C)** frequency of retinaldehyde dehydrogenase (RALDH) positive hepatic DC subsets from naïve (yellow) and 8 week-infected WT mice (blue). **(D)** Representative dot plot staining of FoxP3^+^ Tregs in the liver of CD11c^WT^ and CD11c^ΔAMPKα1^ mice and FoxP3^+^ Tregs as frequency of CD4^+^ T cells. FoxP3^+^ Treg frequencies within CD45^+^ live cells in **(E)** 8 and **(F)** 16 week-infected mice or numbers **(F – right)** in 16 week-infected mice. **(G)** Draining hepatic lymph nodes cells from 8 week-infected mice were restimulated with PMA/Ionomycin in the presence of Brefeldin A and CD4 T cells were analyzed for production of indicated cytokines. **(H)** Draining hepatic and **(I)** mesenteric lymph nodes cells were stimulated with 10μg/mL of soluble egg antigen (SEA) and analyzed for production of indicated cytokines by cytometric bead assay (CBA). **(J)** Frequency of CD11c^-^RELMα^+^ macrophages in the liver of CD11c^WT^ and CD11c^ΔAMPKα1^ mice. **(K)** Representative H&E staining of liver sections (left) and quantification of size of individual egg granuloma (right) from 16 week-infected CD11c^WT^ and CD11c^ΔAMPKα1^ mice. Dashed line marks border of granuloma around the eggs. Bar, 50 μm. Data are from 1 experiment using 3-4 mice per group **(A-C)** or 3-5 mice per group **(D-K).** Student’s T-test **(K)** or Two-Way Anova with Sidák’s post-test **(D-J)** were used to assess statistically significant differences. Mean ± SEM are indicated in the graph.; *p < 0.05, **p < 0.01, ***p < 0.001, ****p<0.0001.

## Discussion

Despite the well-established function for AMPK signaling in promoting catabolic metabolism (Herzig and Shaw, 2017; Garcia and Shaw, 2017) and the growing appreciation for catabolic metabolism as central component for the tolerogenic functions of DCs (Malinarich et al., 2015; Zhao et al., 2018), the importance of AMPK for tolDCs functions has not yet been addressed. Here, we provide evidence that AMPKα1 signaling is crucial for the induction of a tolerogenic phenotype in DCs exposed to RA. Independently from acting on metabolism, AMPKα1 does so by promoting RALDH activity in a FoxO3-dependent manner which is important for the induction of Tregs. In addition, we demonstrate that *in vivo* AMPKα1 signaling in CD11c-expressing cells is important for induction of Tregs during different inflammatory settings.

Characterization of the metabolic properties of the differently generated tolDCs revealed highly divergent profiles, presumably reflecting the differences in molecular targets of Dex, VitD3 and RA through which they modulate DC biology. Most studies interrogating the metabolic properties of human tolDCs have been focused on moDCs treated with VitD3 alone or in conjunction with Dex. Consistent with these earlier reports (Adamik et al., 2022; Ferreira et al., 2015; Vanherwegen et al., 2018; Malinarich et al., 2015), we find VitD3-DCs to display strong metabolic reprogramming with both features of anabolism and catabolism, whereas Dex-DCs showed relatively modest metabolic changes. On the other hand, there have been no detailed investigations into the metabolic properties of RA-DCs. We show here that RA promotes metabolic changes that are distinct from those induced by VitD3 and Dex, such as reduction in ACC1 and Glut1 expression, which may point towards a more catabolic metabolism than the other tolDCs (Vanherwegen et al., 2018). As each of these different tolDCs depend on distinct effector mechanisms to exert their tolerogenic effect, our findings may inform future studies to uncover new functional links between specific metabolic programs/sensors and particular tolerogenic traits of DCs.

The differences in metabolic profiles were also reflected at the level of the key cellular nutrient/energy sensor AMPK, with a potent activation observed in VitD-DCs and RA-DCs, but not in Dex-DCs. This observation would be largely in agreement with earlier reports showing that VitD3, but not Dex treatment, increases mitochondrial biogenesis and OXPHOS (Ferreira et al., 2012), a well-known consequence of increased AMPK signaling (Herzig and Shaw, 2017; Garcia and Shaw, 2017). Indeed, we found that the increase in both OXPHOS and glycolytic rates observed in VitD3-DCs were dependent on AMPK signaling. Surprisingly, however, we found that the AMPK-dependent metabolic reprogramming was dissociated from the tolerogenic functions of VitD3-DCs. Nonetheless, some level of glycolysis and OXPHOS are required for VitD3-DCs to exert their tolerogenic effects, because earlier work has shown that direct inhibition of any of these processes compromises their tolerogenic properties (Ferreira et al., 2015; Vanherwegen et al., 2018).

Interestingly, in contrast to VitD3-DCs, RA-DCs were highly dependent on AMPKα1 signaling to become tolerogenic, both in human and murine *in vitro* models. We found this to be directly linked to the key role that AMPK signaling plays in supporting induction of RALDH activity, a well-known effector mechanism through which RA licenses DCs to promote Treg responses (Bakdash et al., 2015). In addition to RALDH, CD103 expression, another well-established marker induced by RA, was also dependent on AMPK signaling. Interestingly, while AMPK-driven CD103 expression was dependent on PGC1a, we provide evidence for a central role for FoxO3 in mediating AMPK-driven RALDH activity following stimulation with RA. We observed increased phosphorylation levels of AMPK-target site Ser413, (Fasano et al., 2019) which has been shown to increase its transcriptional activity (Greer et al., 2007). While we provide direct support for RA-induced FoxO3 phosphorylation by AMPK at Ser413 in DCs, it is currently still unclear in this context how FoxO3 gets dephosphorylated at Ser253 following RA treatment. mTOR target Akt is known to mediate phosphorylation of FoxO3 at three conserved residues, including Ser253. Because RA has been shown to inhibit mTOR activation (Guo et al., 2021) we can hypothesize that RA treatment of DCs induces AMPK activation leading to an inhibition of Akt-mTOR axis. This would in turn, result in dephosphorylation of FoxO3 at Ser253, increasing its translocation to the nucleus where AMPK-induced phosphorylation of FoxO3 at Ser413 would increase its transcriptional activity. This two-step activation mechanism of FoxO3 by AMPK has been already demonstrated in neurons undergoing excitotoxic apoptosis (Davila et al., 2019) and could also be the mechanism underlying RA signaling in DCs.

How RA triggers AMPK activation, remains to be addressed. Canonically, AMPK gets activated upon cellular bioenergetic stress, reflected by increased ratios of AMP/ATP or ADP/ATP. However, we found no changes in these ratios following RA treatment. Also, inhibition of RAR, the main nuclear receptor through which RA mediates its effects on cells, had no effect on AMPK activity, despite reversing most of RA-induced phenotypic changes in DCs. Interestingly, a recent study revealed that RA may potentially directly interact with AMPK to promote its activity (Zhang et al., 2019) and it is tempting to speculate in the light of our negative findings that this may in fact be the key mode of action through which RA controls AMPK activity in DCs.

We found siLP DCs to have the highest baseline AMPK activation levels amongst DCs from different tissues. Since the intestine is one of the primary sites of RA production, this points towards an important role for RA in promoting AMPK activation in DCs *in vivo*, analogous to our *in vitro* studies. Nonetheless a role for other gut-derived factors in promoting AMPK activation in DCs cannot be excluded. For instance, gut microbiota-derived products such as the short chain fatty acids (SCFA) butyrate, acetate and propionate have been shown to induce AMPK activation in other cell types (Elamin et al., 2013)(Kawaguchi et al., 2002; Sakakibara et al., 2006). Further studies will be needed to define the relative contribution of these local cues to AMPK activation.

The observation that in the siLP AMPK activation status correlated with RALDH activity in DCs and that loss of AMPKα1 resulted in reduced RALDH activity in intestinal CD103^+^CD11b^+^ DCs, as well as in lung CD103^+^DCs, provides direct *in vivo* evidence for an important role of AMPKα1 signaling in underpinning RALDH activity in DCs. While RALDH activity in intestinal and lung CD103^+^ DCs is well known to be linked to the ability of these cells to induce Treg responses (Molenaar et al., 2011; Esterházy et al., 2016, 2019; Denning et al., 2011), CD11c^ΔAMPKα1^ mice did not show clear defects in the intestinal or lung Treg pool under steady state conditions. This may suggest that the reduction in RALDH activity as a consequence of AMPKα1 loss in the DC compartment is not sufficient to affect homeostatic Treg differentiation by these cells. Additionally or alternatively, recent reports suggest that pTreg differentiation at steady state in the gut is mainly controlled by a novel subset of APCs, independently from intestinal conventional DCs (Lyu et al., 2022; Kedmi et al., 2022; Akagbosu et al., 2022). However, of note, these cells are reported to express CD11c, and should therefore also be AMPKα1 deficient in CD11c^ΔAMPKα1^ mice. However, whether that would alter their function remains to be elucidated.

Although we did not find significant changes in Treg populations as a consequence of AMPKα1 deficiency in DCs at steady state, these mice did show a clear defect in Treg accumulation in the context of *S. mansoni* infection as well as HDM-induced allergic asthma. This suggests that AMPKα1 signaling in DCs is required for differentiation and/or expansion during an immunological challenge. In the case of *S. mansoni* infection, this reduction in Treg accumulation was associated with an increased type 2 cytokine response during both acute and chronic stages of the infection and an increase in liver egg-derived granuloma size during the chronic phase of infection. It is well established that during *S. mansoni* infection, Foxp3^+^ Tregs accumulate and that these cells play a crucial role in dampening type 2 immune responses and in keeping the granulomatous response in the liver and gut towards the eggs in check (Turner et al., 2011). Our data would be in line with this concept and suggest that the stronger type 2 immune response and larger granuloma size in infected CD11c^ΔAMPKα1^ are secondary to an impaired Treg response. We should note that, at this point, we cannot formally rule out the possibility that the enhanced type 2 immune response is a direct consequence of loss of AMPKα1 signaling in DCs, for instance by enhancing their capacity to prime Th2 responses. However, this scenario seems unlikely as during HDM challenge, CD11c^ΔAMPKα1^ mice showed a selective defect in the Treg response, without apparent changes in the Type 2 immune response, as determined by eosinophilia. Moreover, others have found that these mice have in fact a compromised type 2 immune response during lung-stage hookworm infection (Nieves et al., 2016). In this infection model we primarily focused on immune responses in the liver. Due to the inherent technical difficulties of reliably obtaining viable single cell suspensions from the gut from infected mice, we were unable to characterize the Treg response in the intestine of these mice but based on the reported similarities in hepatic and intestinal Treg and Type 2 immune responses during this infection (Turner et al., 2011), this is likely to mirror the hepatic immune phenotypes. The appearance of a hepatic CD103^+^CD11b^+^ DC population during *S. mansoni* infection with similarly high AMPK signaling and RALDH activity as their intestinal counterparts would support this. Since hepatic stellate cells in the liver are known to play a central role in RA metabolism and storage and have been shown to release RA upon inflammatory cues (Lee and Jeong, 2012), it is tempting to speculate that during *S. mansoni* infection, this is a direct consequence of increased local exposure of hepatic DCs to RA.

In conclusion, our work has revealed AMPK signaling as a crucial pathway in controlling an RA-induced tolerogenic profile in human CD103^+^ DCs by inducing RALDH activity in a FoxO3-dependent manner, which showed to be functionally important for the induction of Tregs. In addition, we demonstrated that AMPK signaling is key for homeostasis of murine tolerogenic CD103^+^ DCs in the gut and for induction of Treg responses during immunological challenge. Taken together, these data point towards a key role for AMPK signaling in RA-driven development of tolerogenic DCs which may help with the rational design of new therapeutic approaches that target this signaling axis in DCs to control inflammatory responses.

## Methods

### Mice

Itgax-cre Prkaa1-fl/fl (AMPKα1), OVA-specific CD4+ T cell receptor transgenic mice (OT-II) and wild type (WT) mice, both male and female and all on a C57BL/6J background, were bred under SPF conditions at the Leiden University Medical Center (LUMC), Leiden, The Netherlands. Mice were culled through cervical dislocation. Anesthesia with ketamine with either dexdomitor or xylazine was used for *S. mansoni* infection. Animal experiments were performed when the mice were between 8-16 weeks old. Animal experiments were performed in accordance with local government regulations, EU Directive 2010/63EU and Recommendation 2007/526/EC regarding the protection of animals used for experimental and other scientific purposes, as well as approved by the Dutch Central Authority for Scientific Procedures on Animals (CCD). Animal license number AVD116002015253.

### Differentiation of monocyte-derived dendritic cells (moDCs)

moDCs were differentiated from monocytes that were isolated from buffy coats from healthy volunteers that donated blood to the national bloodbank Sanquin (Amsterdam, Netherlands) with informed consent. In brief, blood was diluted 1:2 in Hanks’ Balanced Salt Solution (HBSS) and placed on 12 mL of Ficoll to obtain mononuclear cells. The solution was centrifuged at 600 x g for 30 minutes at 18 ° C, without braking. Subsequently, the formed mononuclear cell layer was removed, placed in a new tube containing HBSS, for further centrifugation at 1000 rpm for 20 min. The cells were washed 2 more times at 1000 rpm for 20 min and monocytes were isolated using CD14 MACS beads (Miltenyi) according to the manufacturer’s recommendations, routinely resulting in a monocyte purity of >95%. Subsequently, 2×10^6^/well of monocytes were cultured in 6 well plate in RPMI medium supplemented with 10% FCS, 100 U/mL of penicillin, 100 μg/mL streptomycin, 2 mM of glutamine, 10 ng/mL human rGM-CSF (Invitrogen) and 0.86ng/mL of human rIL4 (R&D Systems) to differentiate them into moDCs. On day 2/3 of the differentiation process, media was refreshed, and the same concentration of cytokines was added to the cells. On day 5, immature DCs were either treated with 10^-8^ of VitD3 (VitD DCs), 1 μM of retinoic acid (RA DCs), 1 μM of dexamethasone (Dex DCs) or left untreated (mDCs). On day 6, immature DCs were stimulated with 100 ng/mL ultrapure LPS (E. coli 0111 B4 strain, InvivoGen, San Diego). In some experiments, at day 5, moDCs were treated with 10 μM RALDH inhibitor diethylaminobenzaldehyde (DEAB) (Stem Cell Technologies), 10 μM of PPARγ inhibitor (GW9662), 20 μM of PGC1α inhibitor (SR18292) or 1 μM of pan RAR inhibitor (AGN194310) for 30 minutes to 1 hour before treatment with RA.

### Extracellular Flux Analysis

The metabolic characteristics of moDCs were analyzed using a Seahorse XFe96 Extracellular Flux Analyzer (Seahorse Bioscience). In brief, 5×10^4^ moDCs were plated in unbuffered, glucose-free RPMI supplemented with 5% FCS and left to rest 1 hour at 37 °C before the assay in an CO_2_ free incubator. Subsequently extracellular acidification rate (ECAR) and oxygen consumption rate (OCR) were analyzed in response to glucose (10 mM; port A), oligomycin (1 μM; port B), fluoro-carbonyl cyanide phenylhydrazone (FCCP) (3 μM; port C), and rotenone/antimycin A (1/1 μM; port D) (all Sigma-Aldrich). In some experiments, LPS was added in port B after glucose injection.

### RALDH Activity

RALDH activity assay was performed with Aldefluor kit (Stemcell Technologies) according to manufacturer’s protocol. Briefly, cells were harvested, transferred to a V bottom 96 well plate, washed in PBS, stained with Fixable Aqua Dead Cell Stain Kit (Invitrogen) for 15 min at RT and then resuspended in 200 μL of assay buffer. 3 μL (end concentration of 45 μM) of RALDH inhibitor diethylaminobenzaldehyde (DEAB) was added to an empty well. Next, 1 μL (1.83 μM) of Aldefluor reagent was added to the well with the cells and immediately homogenized and transferred to the well containing DEAB. Samples were incubated for 30 min at 37° C and kept in assay buffer until the measurement in a BD FACS Canto. For the *in vivo* experiments, samples were stained with 146 nM of Aldefluor followed by staining with mix of antibodies diluted in assay buffer. Cells were kept in assay buffer until the measurement in a 5 laser Cytek Aurora.

### Human T-cell culture and analysis of T-cell polarization

For analysis of T-cell polarization, moDCs were cultured with allogenic naive CD4^+^ T cells for 7 days in the presence of staphylococcal enterotoxin B (10 pg/mL). On day 7, cells were harvested and transferred to a 24 well plate and cultured in the presence of rhuIL-2 (10 U/mL, R&D System) to expand the T cells. Two days later, 2x concentrated rhuIL-2 was added to the well and cells were divided into 2 wells. Then, 2/3 days later, intracellular cytokine production was analyzed after restimulation with 100 ng/mL phorbol myristate acetate (PMA) and 1μg/mL ionomycin (both Sigma) for a total 6h; 10 μg/mL brefeldin A (Sigma) was added during the last 4 h. Subsequently, the cells were stained with Fixable Aqua Dead Cell Stain Kit (Invitrogen) for 15 min at RT, fixed with 1.9% formaldehyde (Sigma-Aldrich), permeabilized with permeabilization buffer (eBioscience) and stained with anti-IL-4, anti-IL-13, anti-IL-10, anti-IL-17A, anti-IFN-g for 20 min at 4 °C.

### T-cell suppression assay

For analysis of suppression of proliferation of bystander T cells by test T cells, 5×10^4^ moDCs were cocultured with 5×10^5^ human naive CD4^+^ T cells for 6 days. These T cells (test T cells) were harvested, washed, counted, stained with the cell cycle tracking dye 1 μM Cell Trace Violet (Thermo Fisher Scientific) and irradiated (3000 RAD) to prevent expansion. Bystander target T cells (responder T cells), which were allogeneic memory T cells from the same donor as the test T cells, were labeled with 0.5 μM cell tracking dye 5,6-carboxy fluorescein diacetate succinimidyl ester (CFSE). Subsequently, 5×10^4^ test T cells, 2.5×10^4^ responder T cells, and 1×10^3^ allogeneic LPS-stimulated DCs were cocultured for additional 6 days. Proliferation was determined by flow cytometry, by co-staining with CD3, CD4 and CD25 APC. To some cultures, where indicated, 1 μM pan RAR (AGN194310), was added during the DC-T cell coculture.

### Generation of bone marrow-derived GMDCs

BM cells were flushed from mouse femurs and tibia and plated in ‘Nunc™ Cell-Culture Treated 6-well plate’ wells (#140675, Thermo) at a seeding density of 2×10^6^ cells in a volume of 3 mL of complete RPMI (RPMI-1640 supplemented with GlutaMAX™ and 5% FCS, 25 nM β-mercaptoethanol, 100 U/mL penicillin and 100 μg/mL streptomycin) to which 20 ng/mL of recombinant GM-CSF (#315-03, PeproTech, Hamburg, Germany) was added. Cells were differentiated for 8 days with media refreshment on day 4. At day 6, GMDCs were either treated with 1 μM of RA (Sigma) or left untreated. At day 7, GMDCs were either left untreated or activated with 100 ng/mL of LPS (ultrapure, InvivoGen) and after 24 hours semi-adherent cells were harvested for various assays

### In vitro murine T cell priming assay

OT-II cells were negatively isolated with a CD4^+^ T cell Isolation Kit (Miltenyi). 1×10^4^ GMDCs were settled in a round bottom 96-well plate, pulsed with 100 μg/mL of OVA (InvivoGen) in combination with LPS for 5 h, and washed before adding 1×10^5^ CTV-labeled OT-II cells in 200 μL of medium. After 4 days, cells were harvested and analyzed for proliferation by assessing CTV dilution. For cytokine production, cells were left for 6 days and then 100 μL of media was removed and replaced with 100 μL of media containing PMA/Ionomycin (both from Sigma) in the presence of Brefeldin A (Sigma) for 4 hours. Then, cells were stained with Fixable Aqua Dead Cell Stain Kit (Invitrogen) for 15 min at RT, fixed with FoxP3/Transcription Factor Staining Buffer Set (Invitrogen, for FoxP3 detection) for 1 hour at 4 °C, permeabilized with permeabilization buffer (eBioscience) and stained for anti-CD3, anti-CD4, anti-IFN-g, anti-IL-10, anti-IL-17A, anti-FoxP3 and anti-CD25.

### Flow cytometry

In general, single cell suspensions underwent viability staining for 20 minutes at room temperature (RT) using the LIVE/DEAD™ Fixable Aqua Dead Cell Stain Kit (#L34957, Thermo; 1:400 in PBS [from LUMC pharmacy; Braun, Zeist, The Netherlands) or LIVE/DEAD™ Fixable Blue Cells Stain Kit (#L34957, Thermo; 1:1000 in PBS) and fixation for 15 minutes at RT using 1.85% formaldehyde (F1635, Sigma) in PBS solution before surface staining with antibodies in PBS supplemented with 0.5% BSA [fraction V, #10735086001, Roche, Woerden, The Netherlands] and 2 mM EDTA) for 30 minutes at 4 °C. For detection of metabolic proteins cells were permeabilized with eBioscience permeabilization buffer (#00-8333-56 – Thermo) followed by intracellular staining with a cocktail of antibodies against the metabolic proteins. Antibodies were purchased from AbCam and conjugated in house with AbCam Lightening-Link Conjugation kit. See supplementary table 1 for further information on metabolic antibodies and fluorochrome conjugation. For polyclonal restimulation, single cell suspensions were stimulated with both 0.1 μg/mL of PMA (#P-8139, Sigma) and 1 μg/mL of ionomycin (#I-0634, Sigma) in the presence of 10 μg/mL BrefeldinA (all Sigma Aldrich) for 4 hrs. These single cell suspensions underwent intracellular cytokine staining (ICS) with antibodies in 1x eBioscience™ Permeabilization Buffer (ebioscience - # 00-8333-56).

For detection of phosphorylated ACC on Ser79, cells were pre stained with LIVE/DEAD™ Blue Dead Cell Stain Kit for 20 min at RT and, after washing, single cell suspension was returned to a cell incubator (37°C & 5% CO2) for 1 hour in cRPMI and fixed with 4% ultra-pure formaldehyde (#18814-20, Polysciences, Hirschberg an der Bergstraβe, Germany) for 15 minutes at RT. Single cell suspensions were first stained with anti-phosphorylated ACC in 1x Permeabilization Buffer (#00-8333-56, Thermo) for 1 hour at room before staining with cocktail antibodies in 1x Permeabilization Buffer for 30 minutes at 4°C.

### High dimensional spectral flow cytometry analysis

Samples were imported in OMIQ software and parameters were scaled using a conversion factors ranging from 6000-20000. Samples were subsequently gated on live moDCs or CD64^-^CD11c^+^MHCII^+^ intestinal DCs and subsampled using a maximum equal distribution across groups. After sub-sampling, UMAP analysis was performed using the metabolic proteins (in case of moDCs) or lineage markers and activation markers (in case of intestinal DCs) as parameters. Next, phenograph clustering (k=100) was performed using the same parameters used for the UMAP. Data was further analyzed with EdgeR to determine significant differences in the clusters among different genotypes and heatmaps and volcano plots were generated in R, using OMIQ-exported data for each cluster.

### Quantitative real-time PCR

RNA was extracted from snap-frozen moDCs. The isolation of mRNA was performed using Tripure method. Briefly, 500 μL of Tripure reagent (Sigma) and 1 μL of RNAse-free glycogen (Invitrogen) were added to the pellet of samples. Samples were then vortexed and incubated at RT for 5 min. After this step, 100 μL of RNAse-free chlorophorm was added and samples were inverted vigorously for 30 seconds. Samples were then centrifuged for 12000 x g for 15 min at 4 °C and the upper aqueous phase was transferred to a new RNAse-free 1.5 mL tube. Then, 250 μL of 2-isopropanol was added to each tube and after 5 min of incubation samples were centrifuged for 12000 x g for 15 min at 4 °C. Next, 500 μL of 75% ethanol was added, tubes were vortexed and centrifuged for 12000 x g for 15 min at 4 °C. As much as possible of ethanol was removed and 30 μL of RNAse-free milliQ water was added to the tubes that were incubated for 10 minutes at 55 °C. After this, RNA samples were quantified with NanoDrop (Invitrogen). cDNA was synthesized with reverse transcriptase kit (Promega) using the same amount of RNA for all the samples (settled by the sample with the lowest concentration of RNA). PCR amplification was performed by the SYBR Green method using CFX (Biorad). GAPDH mRNA levels was used as internal control. Specific primers for detected genes are listed in the supplementary Table 2. Relative expression was determined using the 2^ΔΔCt^ method.

### Small interfering RNA (siRNA) electroporation

On day 4 of the DC culture, the cells were harvested and transfected with either 20 nM non-target control siRNA or 20 nM AMPKα1 (both Dharmacon) or FoxO3 (Invitrogen) siRNA using Neon Transfection System (Invitrogen) with the following setting: 1.600 V, 20 ms width, one pulse. Following electroporation, 1.2×10^6^ cells were seeded per well in to a 6-well plate containing RPMI media without antibiotics. After 24 h, culture medium (RPMI) supplemented with 10% FCS, rIL-4 (0.86 ng/mL, R&D system) and rGM-CSF (10 ng/mL, Invitrogen) was added. AMPKα1 and FoxO3 silencing efficiency was determined by qPCR and the transfection efficiency was greater than 90%.

### Western blotting

A minimum of one million GMDCs or moDCs were washed twice with PBS before being lysed in 150 μL of EBSB buffer (8% [w/v] glycerol, 3% [w/v] SDS and 100 mM Tris–HCl [pH 6.8]). Lysates were immediately boiled for 5 min and their protein content was determined using a BCA kit. Ten μg of protein per lain was separated by SDS-PAGE followed by transfer to a PVDF membrane. Membranes were blocked for 1 h at room temperature in TTBS buffer (20 mM Tris–HCl [pH 7.6], 137 mM NaCl, and 0.25% [v/v] Tween 20) containing 5% (w/v) fat free milk and incubated overnight with primary antibodies. The primary antibodies used were AMPK (Cell Signaling #2532), pAMPK Ser79 (Cell Signaling #2535S), FoxO3 (Cell Signaling #2497S), pFoxO3 Ser413 (Cell Signaling #8174S), pFoxO3 Ser 253 (Cell Signaling #9466), HSP90 (Santa Cruz #sc7947) and beta-actin (Sigma #A5441). The membranes were then washed in TTBS buffer and incubated with horseradish peroxidase-conjugated secondary antibodies for 2 hours at RT. After washing, blots were developed using enhanced chemiluminescence.

### Digestion of mouse tissues

Lymphoid organs were collected in 500 μL of no additives media (naRPMI = RPMI-1640 supplemented with GlutaMAX™ [#61870-010 or alternatively 61870036, Gibco, Bleiswijk, The Netherlands], in a plate and mechanically disrupted using the back-end of a syringe before addition of 50 μL of a digestion media (dRPMI = naRPMI supplemented with 11x collagenase D (#11088866001, Roche, Woerden, The Netherlands; end concentration of 1 mg/mL) and 11x DNase I (#D4263, Sigma, Zwijndrecht, The Netherlands; end concentration of 2000 U/mL) for 20 minutes at 37°C and 5% CO2. Single cell suspensions were filtered after digestion with a 100 μm sterile filter (#352360, BD Biosciences, Vianen, The Netherlands) before counting in complete RMPI (cRPMI = naRPMI supplemented with 10% heat-inactivated FCS [#S-FBS-EU-015, Serana, Pessin, Germany], 25 nM β-mercaptoethanol [#M6250, Sigma], 100 U/mL penicillin [#16128286, Eureco-pharma, Ridderkerk, The Netherlands; purchased inside the LUMC] and 100 μg/mL streptomycin [#S9137, Sigma]). Spleens were subjected to red blood cell lysis (inhouse; 0.15 M NH4Cl, 1 mM KHCO3, 0.1 mM EDTA [#15575-038, Thermo, Waltham, Massachusetts, United States] in ddH20) for 2 minutes at room temperature before counting.

### Small intestine lamina propria isolation

Intestinal LP cells were isolated, as previously described(Scott et al., 2016). Briefly, intestine was collected, and fat tissues and payer’s patch were manually removed. Small intestine was open longitudinally and feces and mucus were removed using PBS and tissues were cut in small pieces of approximately 2 cm and transfer to a 50 mL tube with HBSS no Ca^2+^ or Mg^2+^ (Gibco) supplemented with 10% FCS and 2 mM EDTA. The pieces of intestine were manually shook for 30 seconds, poured into a nitex nylon mesh, and washed again with plain 10 mL of HBSS no Ca^2+^ or Mg^2+^ (Gibco). Tissues were then transferred back to 50 mL tubes, now containing 10 mL of pre warmed HBSS no Ca^2+^ or Mg^2+^ (Gibco) + 2 mM EDTA and placed in an orbital shaker at 200 rpm for 20 min at 37 °C. This procedure was repeated three more times and after the third washing, guts were transferred to a new 50 mL tubes containing 1x digestion mix (RPMI GlutaMAX™ containing 2% FCS, 1mg/mL of Colagenase VIII [Sigma] and 40 u/mL of DNAse I [Sigma]) and incubated for 20 min at 37 °C sin an orbital shaker at 200 rpm. After digestion, 35 mL of PBS containing 2% FCS and 2 mM EDTA was added to the samples and placed in a 100 μm cell strainer on top of a new 50 mL tube. Samples were centrifuged at 450 x g for 5 min, resuspended in 10 mL of PBS containing 2% FCS and 2 mM EDTA, place in a 40 μm cell strainer and centrifuged at 450 x g for 5 minutes.

### *Schistosoma mansoni* acute and chronic infection

Mice were infected with S. mansoni (Puerto Rican strain; Naval Medical Research Institute) by 30 minutes of percutaneous exposure to 35 cercariae on shaved abdomen. Mice were culled 8 weeks later. Cercariae were kept at 30 cercariae per mL of store bought Barleduc water, which was kept very carefully at 31 degrees Celsius. Female mice were anesthetized by intraperitoneal (i.p) injection with 300 μL of 50 mg/kg bodyweight ketamine + 0.5 mg/kg bodyweight dexdomitor, while male mice were anesthetized with 50 mg/kg bodyweight ketamine + 10 mg/kg bodyweight xylazine. Female mice were assisted in waking up by i.p. injection with 150 μL of 0.4 mg/kg bodyweight. All injections were done using PBS and a 25G needle. All anesthetics were purchased at the LUMC pharmacy. Livers were processed like spleens except that single cell suspensions were centrifuged twice at 20 g for 10 minutes in PBS to remove hepatocytes before red blood cell lysis.

### House dust mite-induced allergic asthma model

Allergic airway inflammation was induced by sensitizing mice via intranasal administration of 1 μg HDM (Greer, London, United Kingdom) in 50 μL of PBS. One week later, these mice were challenged for 5 consecutive days via intranasal administration of 10 μg HDM in 50 μL of PBS. On day 15, bronchoalveolar lavage (BAL) fluid, lung, and lung draining mediastinal LNs (med LNs) were obtained to determine inflammatory cell recruitment. BAL was performed by instilling the lungs with 3 × 1mL aliquots of sterile PBS (Braun, Oss, The Netherlands). For single-cell suspensions of whole lung tissue, lungs were perfused with sterile PBS via the right ventricle to clear leukocytes and erythrocytes from the pulmonary circulation. Lung and medLN homogenization were performed as described above. Eosinophilia was assessed in BAL and lungs by flow cytometry as a readout for allergic inflammation.

### Cytometric bead array

Cell culture supernatants were analyzed for IFNγ, IL-4, IL-5, IL-10 and IL-13 secretion using a cytokine bead array (#I558296, #558298, #558302, #558300 and #558349 respectively and all BD Biosciences) on a flow cytometer as recommended by the manufacturer, but with both the beads and antibodies diluted 1:10 relative to the original recommendation.

### Adenine nucleotides concentration

The intracellular concentration of adenosine mono, di or triphosphate (AMP, ADP, ATP, respectively) were evaluated as previously described (García-Tardón and Guigas, 2018). Briefly, cells were harvested, transferred to a 1.5 mL tube and washed in ice-cold PBS. After centrifugation cells were rinsed with 250 μl of perchloric acid/EDTA [10% (v/v) HClO4, 25 mM EDTA] and, vortexed and kept on ice for 30 minutes. Then, cell extracts were spun down at 8000 x g for 2 minutes at 4 °C and 200 μl of the supernatant fraction was transferred into a new 1.5mL tube (on ice), where cells lysate was neutralized by adjusting the pH to 6.5-7 with approximately 130 μl of KOH/-MOPS [2 N KOH/0.3 M 3-(N-morpholino)propanesulfonic acid (MOPS)]. Lysate was centrifuged at 8000 x g for 2 minutes at 4 °C and 200 μl of the neutralized fraction was transferred to a new 1.5 mL tube and stored in −80 °C until measurement of AMP, ADP and ATP at ultraviolet (UV)-based high-performance liquid chromatography (HPLC - DIONEX UltiMate 3000, Thermo Scientific).

### Statistical analysis

Results are expressed as mean ± standard error mean (SEM) except stated otherwise. Continuous variables were log-transformed for the analyses when the normality of the distribution was rejected by the Shapiro-Wilk W test and in case of a new normality test failure, the nonparametric alternative was used for the analysis. Differences between groups were analyzed by two-way ANOVA or Kruskall-Wallis (non-parametric analysis) with Tukey HSD or Sidak’s multiple comparison post-test as described in figure legends. If there were only two groups, unpaired Student’s t-test or Mann-Whitney test (non-parametric analysis) were used. *p* values < 0.05 were considered significant (*p < 0.05, **p < 0.01, ***p < 0.001) and statistical analyzes were performed using GraphPad Prism v.9.2.

### Online supplemental material

**Figure S1** shows the metabolic and phenotype characterization of tolerogenic human moDCs. **Figure S2** shows the ECAR and OCR from Seahorse extracellular flux analysis of immunogenic and tolerogenic DCs after real-time LPS restimulation. **Figure S3** shows the characterization of AMPKα1 deletion in both human and murine DCs and the metabolic characterization of Dex-DCs silenced for AMPKα1. **Figure S4** shows the phenotypic characterization of RA-DCs silenced for either AMPKα1 or FoxO3 and the T cells priming capacity of both GMDCs treated with RALDH inhibitor (DEAB) and moDCs treated with either PPARγ (GW9662) or PGC1α (SR18292) inhibitors. **Figure S5** shows the gating strategy to identify different murine DC subset sand macrophages within tissues. **Figure S6** shows AMPK signaling in different DC subsets within tissues and the characterization of migratory DCs and regulatory T cells in the msLN of CD11c^WT^ and CD11c^ΔAMPKα1^ mice. **Figure S7** shows the characterization of lungs from CD11c^WT^ and CD11c^ΔAMPKα1^ mice under steady state condition or after house dust mite challenge.

## Acknowledgments

This work was supported by an LUMC fellowship and NWO ENW grant (OCENW.M.21.057) awarded to BE. The authors want to acknowledge the São Paulo Foundation Research for a fellowship awarded to TAP (# 2018/00719-9 and #2014/26437-9).

## Author Contribution

TAP and BE designed the experiments, interpreted data. TAP, ECB, GH, LP, AZD, FO, AOF, AJvH and BG performed the experiments. JAMB helped in the discussion and reviewed the manuscript. BE conceived and supervised the study and wrote the manuscript together with TAP.

