## Supplementary figures and images for "Metabolic sensor AMPK licenses CD103^+^ dendritic cells to induce Treg responses"

### Figure Supplementary 1.pdf

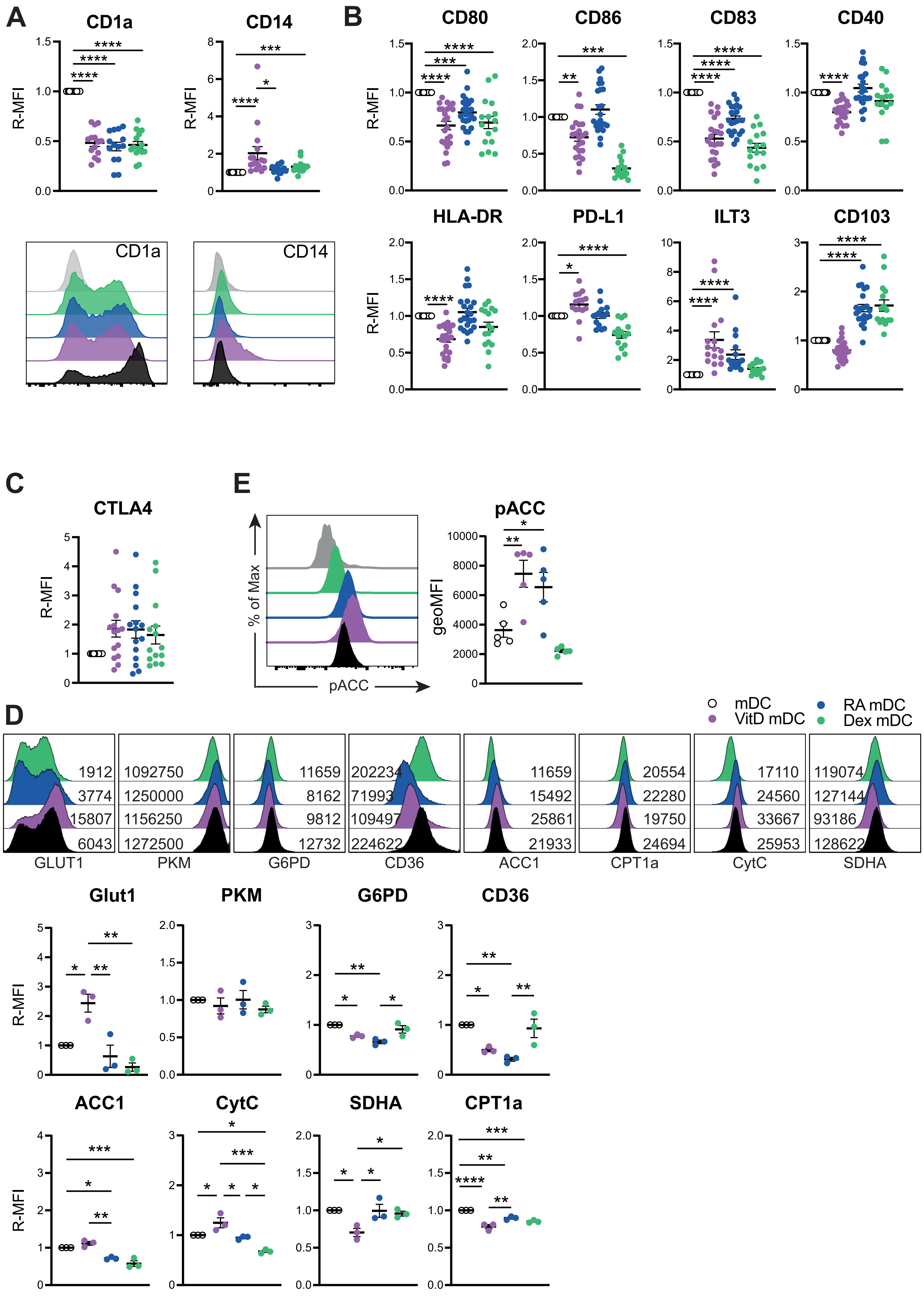

### Figure Supplementary 2.pdf

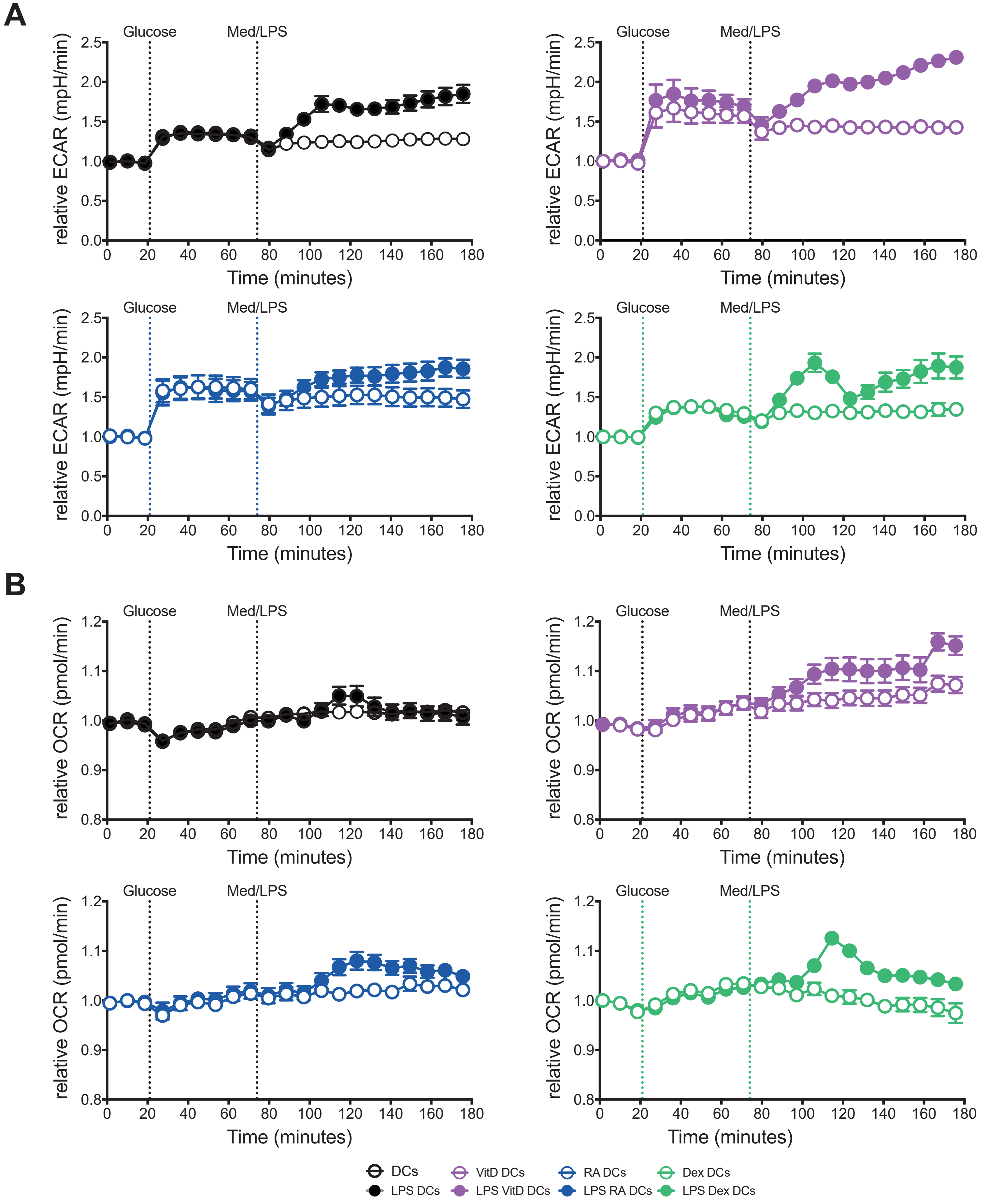

### Figure Supplementary 3.pdf

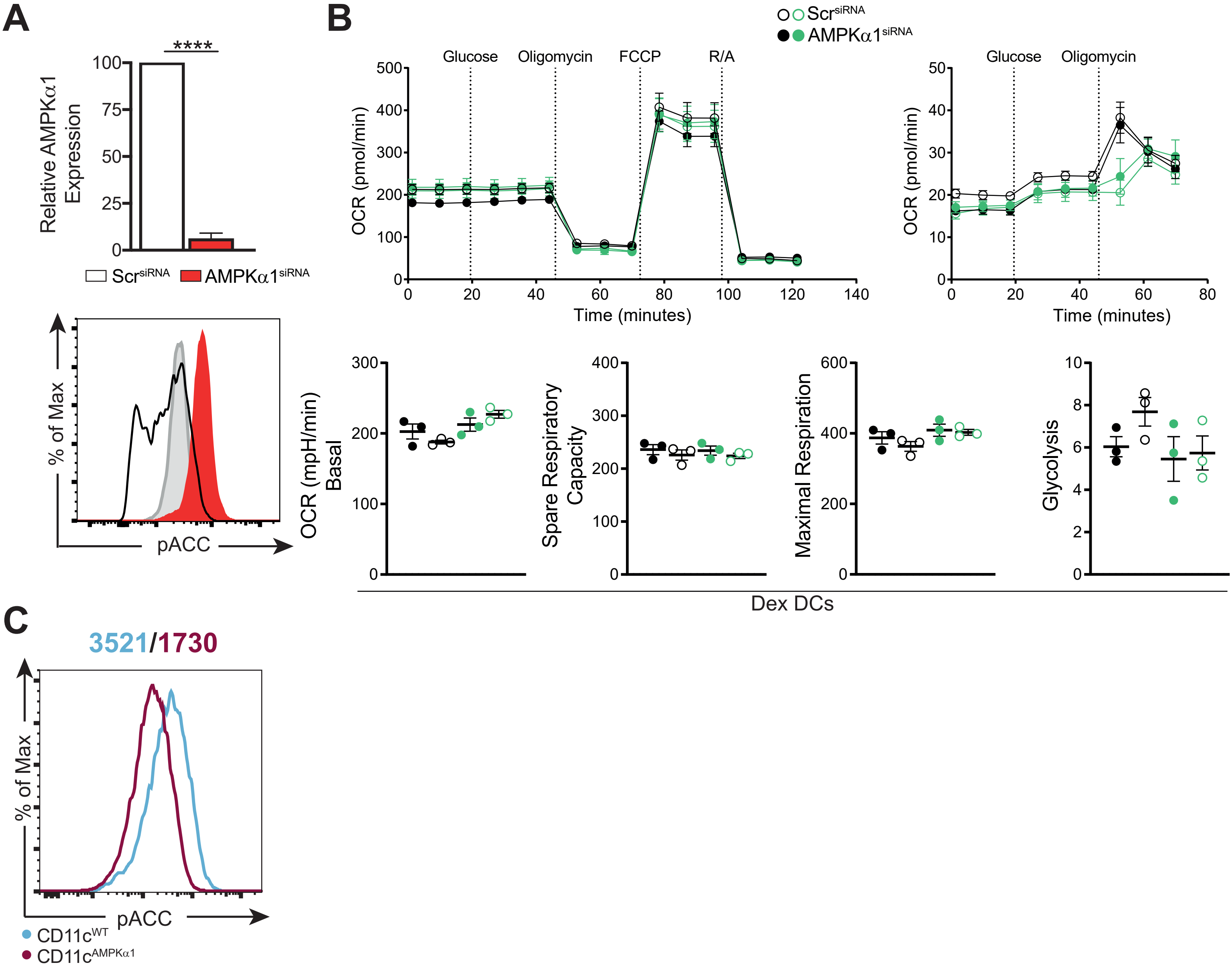

### Figure Supplementary 4.pdf

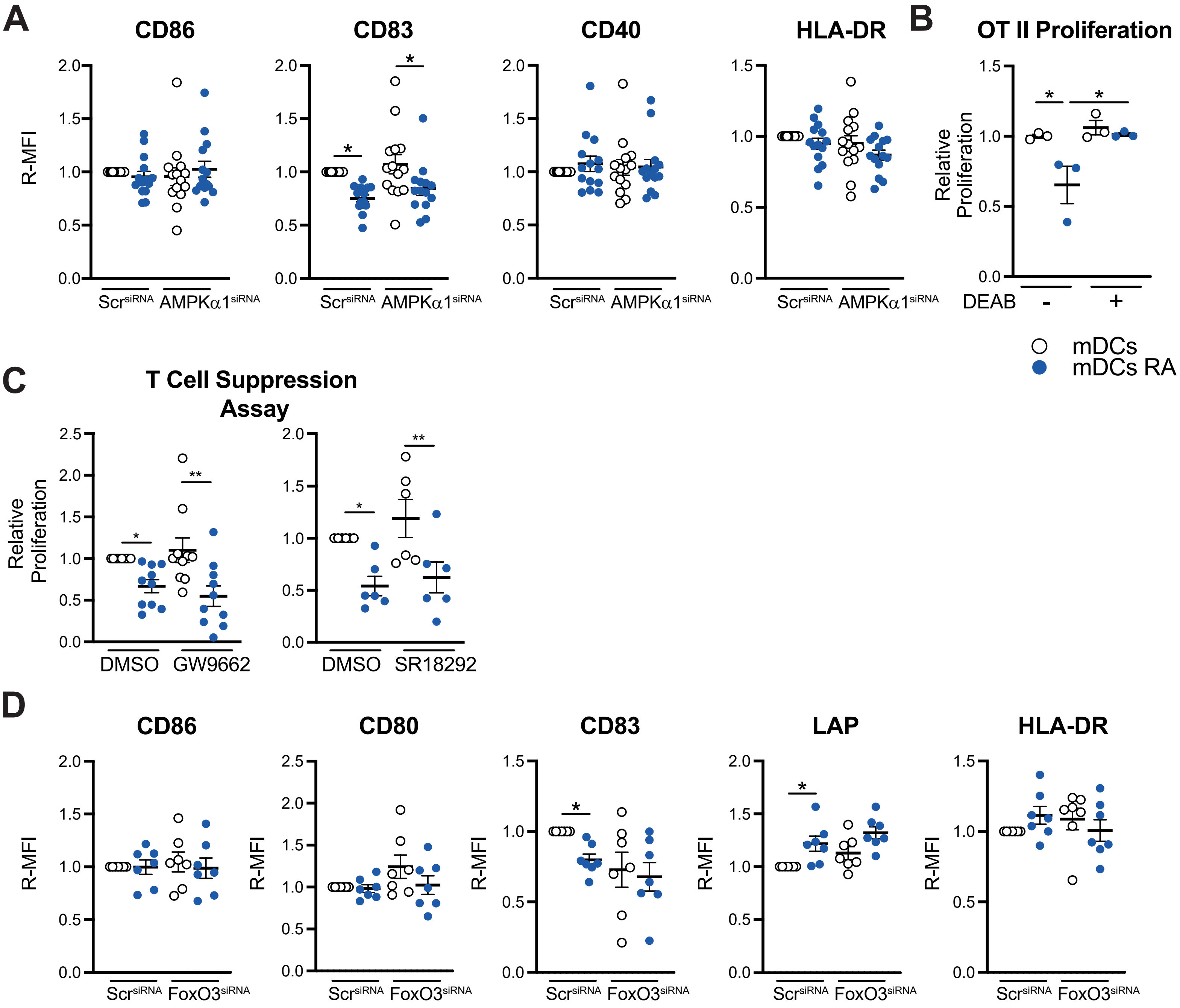

### Figure Supplementary 5.pdf

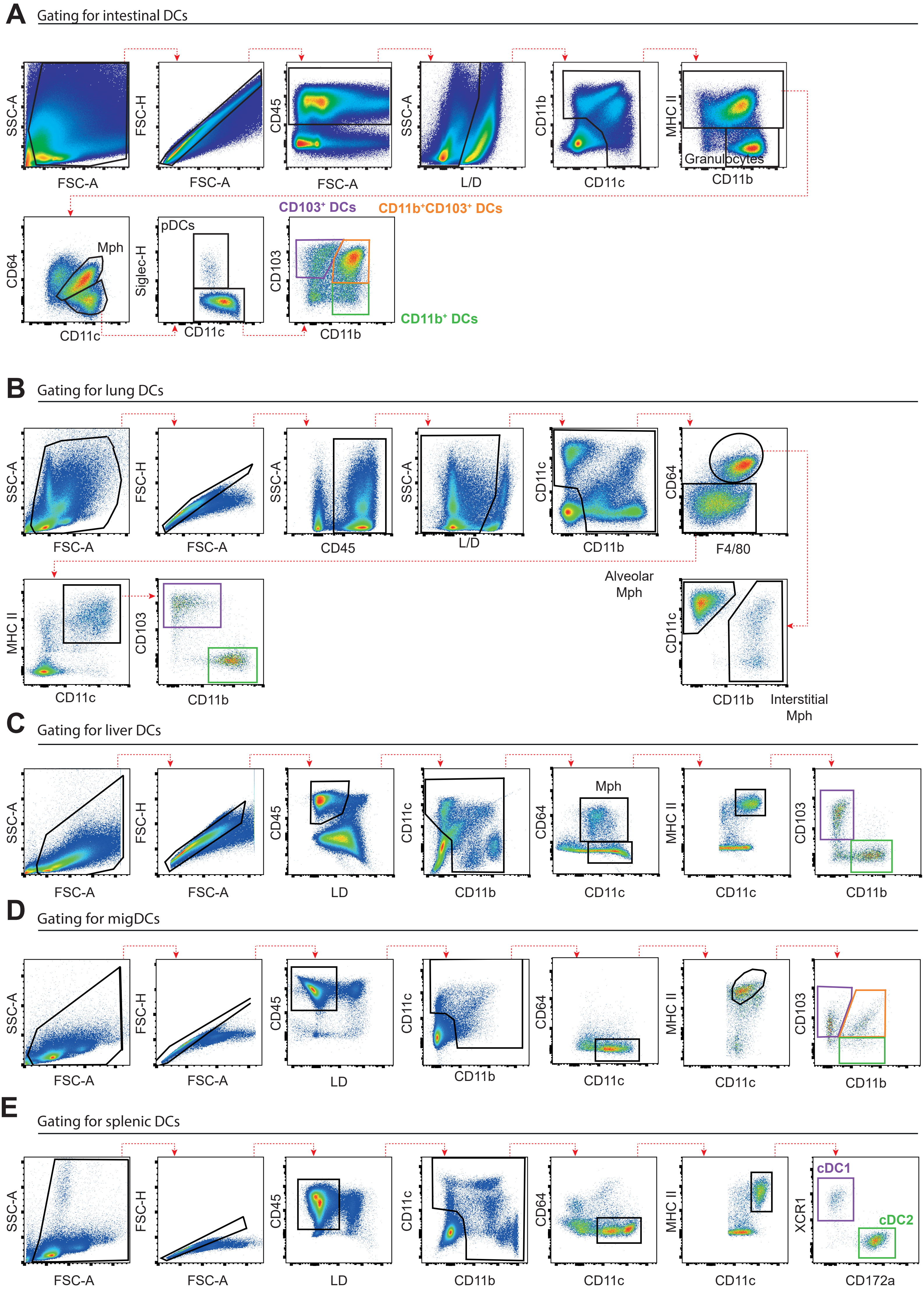

### Figure Supplementary 6.pdf

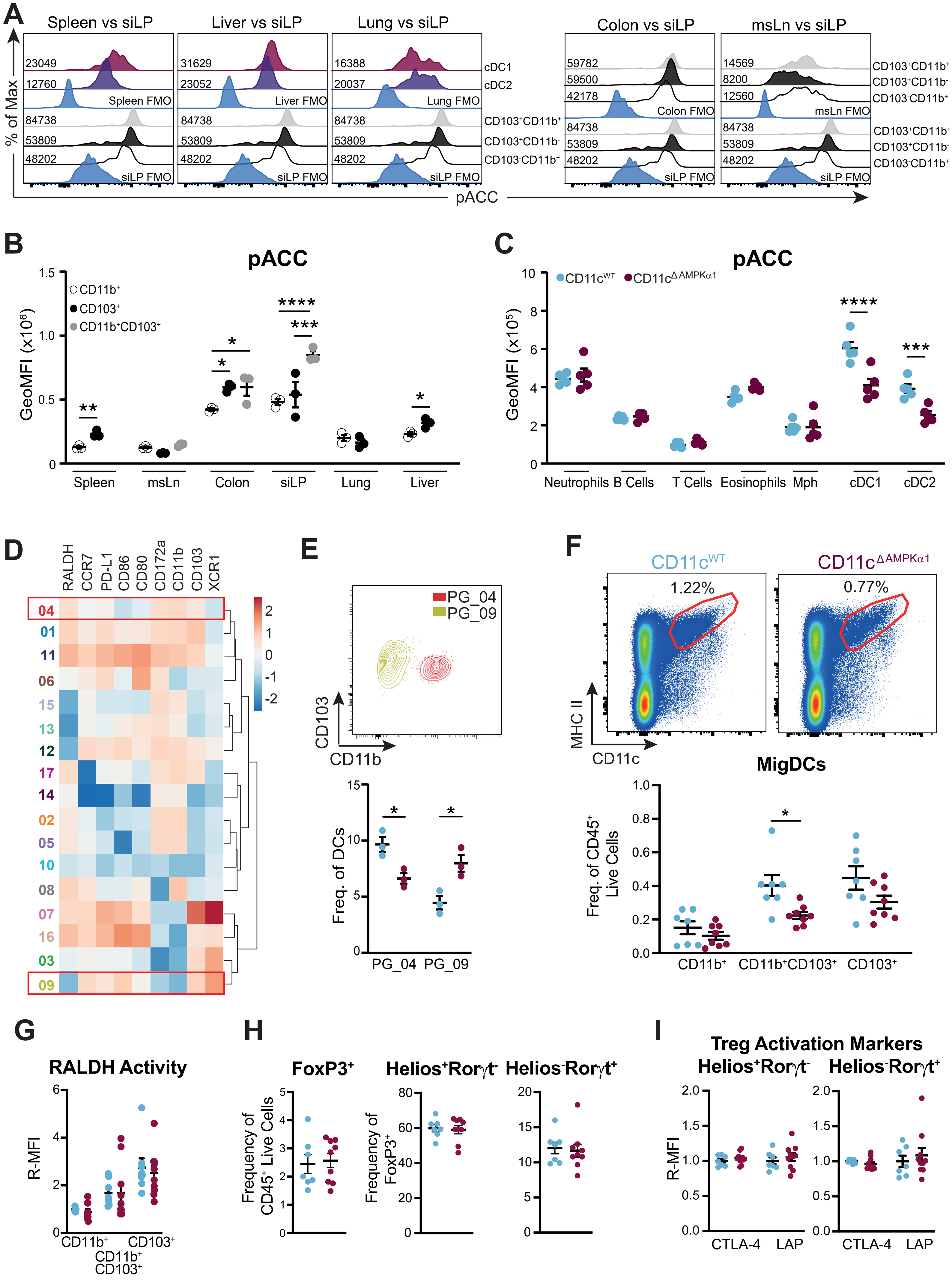

### Figure Supplementary 7.pdf

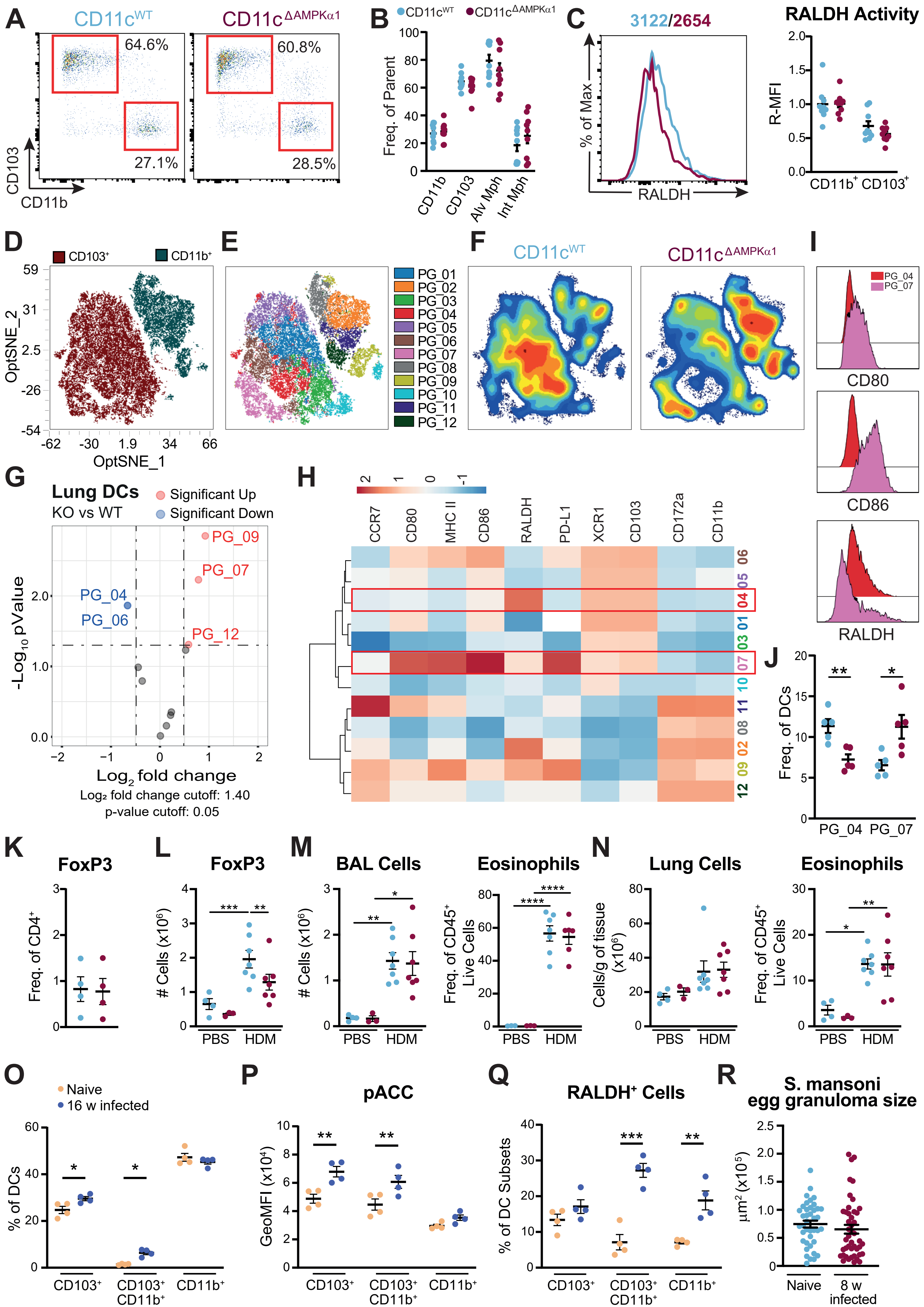
